# Nanoscale imaging reveals the mechanisms of ER-to-Golgi transport via a dynamic tubular-vesicular network

**DOI:** 10.1101/2023.10.27.563951

**Authors:** Luis Wong-Dilworth, Gresy Bregu, Steffen Restel, Carmen Rodilla-Ramirez, Svenja Ebeling, Shelly Harel, Paula Leupold, Jonathan Grimm, Luke D. Lavis, Jessica Angulo-Capel, Felix Campelo, Francesca Bottanelli

**Author notes:** These authors contributed equally (Gresy Bregu, Steffen Restel, Francesca Bottanelli).

## Abstract

The endoplasmic reticulum (ER) and the Golgi apparatus are the first sorting stations along the secretory pathway of mammalian cells and have a crucial role in protein quality control and cellular homeostasis. While machinery components mediating ER-to-Golgi transport have been mapped, it is unclear how exchange between the two closely juxtaposed organelles is coordinated in living cells. Here, using gene editing to tag machinery components, live-cell confocal and stimulated emission depletion (STED) super-resolution microscopy, we show that ER-to-Golgi transport occurs via a dynamic network of tubules positive for the small GTPase ARF4. swCOPI machinery is tightly associated to this network and moves with tubular-vesicular structures. Strikingly, the ARF4 network appears to be continuous with the ER and ARF4 tubules remodel around static ER exit sites (ERES) defined by COPII machinery. We were further able to dissect the steps of ER-to-Golgi transport with functional trafficking assays. A wave of cargo released from the ER percolates through peripheral and Golgi-tethered ARF4 structures before filling the cis-Golgi. Perturbation via acute degradation of ARF4 shows an active regulatory role for the GTPase and COPI in anterograde transport. Our data supports a model in which anterograde ER-to-Golgi transport occurs via an ARF4 tubular-vesicular network directly connecting the ER and Golgi-associated pre-cisternae.

## Introduction

Cells compartmentalize biochemical reactions into interconnected membrane bound organelles. The endoplasmic reticulum (ER) is the first organelle along the endomembrane system and has a crucial role in cellular quality control as it is involved in folding and assembly of 1/3 of the total cellular proteins^1^. Fully folded proteins are then sorted for transport to the Golgi apparatus by accumulating at ER sub-domains called ER exit sites (ERES)^2^. The process of cargo selection and packing is driven by COPII machinery^3^. According to established textbook models, homotypic fusion of budding COPII vesicular carriers at ERES would lead to the formation of the ER-Golgi intermediate compartment (ERGIC), a transient sorting station at the interface between ER and the first cis-Golgi cisterna which receives incoming cargo^4^. Early imaging of ERGIC dynamics suggested various possible models as to how cargoes are sorted at the level of the intermediate compartment in an anterograde (ERGIC-to-Golgi) or retrograde (ERGIC-to-ER) manner^5^. COPI machinery, whose role in cis-Golgi-to-ER retrograde transport is well established, could additionally lead to the formation of Golgi-destined COPI vesicles from static ERGIC membranes. Another possibility is that the entire ERGIC would translocate to deliver cargo to the Golgi. Up to date, all these models remain untested and the exact roles of COPI in bi-directional transport at the ERGIC are ill defined. In support for a role of COPI in anterograde traffic, antibodies against the β subunit of the COPI coat disrupted anterograde traffic of the model cargo vesicular stomatitis virus G protein (VSV-G) which was arrested in the ERGIC^6^. Additionally, dynamic Golgi-bound COPI-positive structures have been visualized next to long-lived and static COPII punctae, presumably representing ERES^7,8^. Recent publications have challenged the classical vesicular model and have proposed that COPII machinery localizes to the neck of a tubular-vesicular network that serves as a cargo export site^9,10^. This interwoven tubular-vesicular network itself, rather than vesicles, would be responsible for anterograde cargo export in mammalian cells via continuities with the ER lumen^10^. It has been speculated that COPII machinery would select and shuffle cargoes into tubular-vesicular structures and, at the same time, provide a diffusion barrier to avoid lipid and protein mixing and help retaining the identity of the two organelles^10^. Machinery like TANGO1, a protein that is able to simultaneously bind cargo and the COPII coat, could support the formation of tunnels and transient connections between the ER and downstream compartments^11^. TANGO1 would prevent severing of a vesicle and promote membrane expansion at ER export sites by recruiting tethers for the recruitment of ERGIC membranes^11–13^.

ARF GTPases are key regulators of intracellular membrane communication and have a key role in COPI recruitment and early secretory traffic^14^. They are divided in 3 classes based on sequence similarity and are all strikingly structurally homologous. Type I ARF1 and Type II ARF4 and ARF5 all support COPI vesicle formation *in vitro*^15^ and analysis of their nanoscale localization with stimulated emission depletion (STED) super-resolution microscopy has recently revealed proximity to different COPI sub-populations on the ERGIC (ARF4 and ARF5) and the first cis-Golgi cisterna (ARF1)^16^. While the best studied of the ARF paralogues, ARF1, which has a role in retrograde Golgi-to-ER transport^17^, no ARF GTPase has been so far attributed a role in the regulation of ER-to-Golgi anterograde transport. Intriguingly, ARF1 and ARF4 localize to different populations of ERGICs^16^. ARF4 defines a population of ERGICs tightly tethered to the first cis-Golgi cisternae, which we have termed Golgi-associated ERGICs (GA-ERGICs). ARF4 also localizes to peripheral ERGICs (p-ERGICs) disperse throughout the cytoplasm. ARF1, however, only localizes to a sub-set of p-ERGICs and is excluded from GA-ERGICs, suggesting that functionally different sub-populations of ERGICs may co-exist. Although the function of ARF4 in trafficking is still elusive^18^, a role in formation of tubular carriers from the ERGIC has been suggested as simultaneous knock-down of both, ARF1 and ARF4, causes extensive tubulation of ERGIC membranes^19,20^. Interestingly, an ARF4 knock-out (KO) led to the secretion of ER resident proteins, suggesting a role in retrieval from ERGICs^21^.

Here we use fast confocal and live-cell STED super-resolution microscopy^22^ to follow the dynamics of gene-edited ARFs, COPI and COPII to dissect steps of ER-Golgi transport in human cells. We show that ARF4 defines a highly pleiomorphic and dynamic network of tubules connecting the ER and GA-ERGICs. COPI is tightly associated to this network and moves with tubular-vesicular structures. Strikingly, the ARF4 tubular network appears to be continuous with and tightly associated to the ER and remodels around static ERES defined by COPII machinery. A wave of anterograde cargo is first observed within ARF4 tubules and GA-ERGICs before filling the first cis-Golgi cisterna. Acute depletion of ARF4 triggers release of COPI from membranes and mis-regulation of ER-Golgi transport, suggesting a direct role for ARF4 and COPI in tuning of anterograde cargo flow.

## Results

To dissect steps of ER-Golgi transport, we followed the dynamics of gene-edited ARFs, COPI and COPII components using fast confocal and live-cell STED super-resolution microscopy in HeLa and RPE-1 cells human cell culture models. Fast-spinning disk confocal microscopy of endogenously Halo-tagged ARF4 cells (ARF4^EN^-Halo, EN=endogenous) shows a highly pleiomorphic and quickly remodeling tubular-vesicular network and peri-nuclear signal (Figure 1A, Supplementary Movie 1). As previously shown, ARF4-positive structures are positive for the ERGIC marker ERGIC53 and are therefore intermediate compartments sitting at the interface between the ER and the Golgi making us wonder what their role in ER-ERGIC-Golgi transport is (Extended Data Fig. 1A,C)^16^. Importantly, the same localization for edited ARF4 on GA-ERGICs is observed in non-cancerous RPE-1 cells (Extended Data Fig. 1B,D). Live-cell STED microscopy is able to discern in both cell types different sub-classes of ARF4-positive ERGICs that are either tethered to the cis-Golgi cisternae (GA-ERGICs) (Extended Data Fig. 1Ci-Di) or in the cell periphery (p-ERGICs) (Extended Data Fig. 1Cii-Dii). Again, live-cell STED shows that ARF4 GA-ERGICs are tethered to the cis-Golgi cisternae defined by gene edited ARF1 (ARF1^EN^-SNAP) in both HeLa and RPE-1 cells (Extended Data Fig. 2A-B). The ARF4 network is characterized by tubules that 1) bridge p-ERGICs (Fig. 1A-Ia), 2) connect p-ERGICs with GA-ERGICs (Fig. 1A-Ib) and 3) translocate from the periphery towards the Golgi to become GA-ERGICs (Fig. 1A-II and Fig. 1B and H). ARF4 tubules are mostly segregated from ARF1 tubules, which mediate Golgi-to-ER recycling or post-Golgi trafficking^17^. Interestingly, tubules of possible retrograde (ARF1) and anterograde (ARF4) nature could form from the same ERGIC (Extended Data Fig. 2C-D, Supplementary Movie 2). Further, ARF4 forms highly mobile tubules that emanate from static p-ERGICs (Fig. 1C-D, Extended Data Fig. 3). To better characterize the dynamics, we tracked the movement of ARF4 structures and use the aspect ratio as a metric to distinguish between elongated (highly mobile tubules) and rounded structures (static p-ERGICs) (Extended Data Fig. 3A-C). Accordingly, we observed that elongated structures (aspect ratio > 1.6) exhibit significantly higher instantaneous velocities and diffusion coefficients than rounded structures (aspect ratio < 1.6) (Fig. 1C-D, Extended Data Fig. 3B-E). To further dissect the localization, dynamics and function of COPI machinery on ARF4 ERGIC compartments, we took advantage of a βCOP (a subunit of the COPI coat) and ARF4 double KI cell line^16^. Live-cell STED imaging of endogenous ARF4 and βCOP shows that ARF4 tubules are defined by nanodomains of COPI (Fig. 1E) and fast dynamic imaging highlights that COPI remains associated to translocating ARF4-positive tubular-vesicular structures rather than budding vesicles (Fig. 1F, Supplementary Movie 3). Dynamic live-cell STED imaging captures tubular-vesicular ARF4-COPI structures establishing tubular connections where COPI is localized to the connecting tubular bridge (Fig. 1G Ia and Ib and Fig.1H) and an incoming tubular-vesicular structure becoming tethered to the Golgi cisterna (Fig. 1G II and Fig. 1H, Supplementary Movie 4). Our data shows that ARF4 defines a dynamic ERGIC network with stably associated COPI nanodomains.

**Figure 1.**
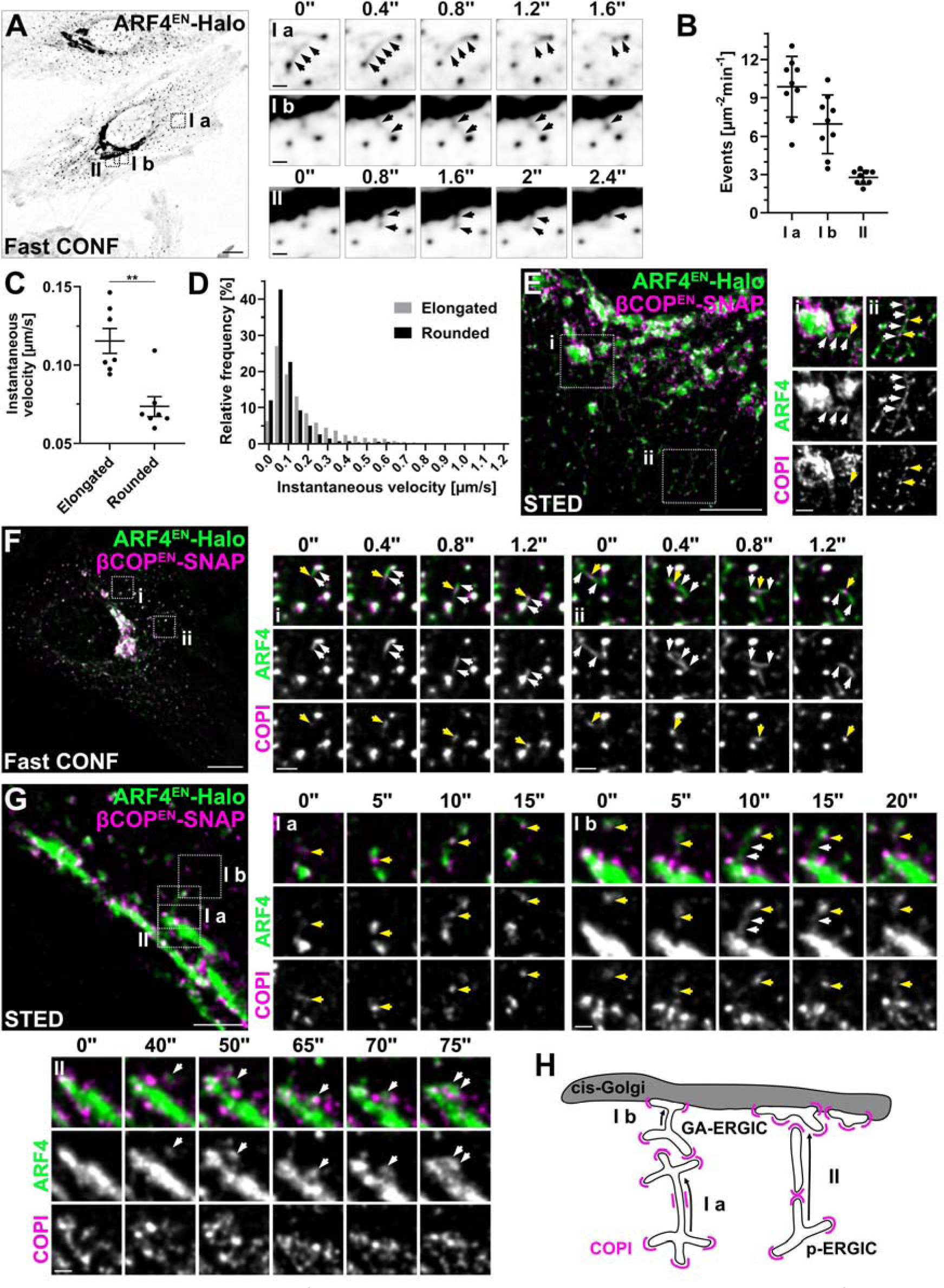
ARF4 and COPI define a highly dynamic tubular-vesicular network of ER-Golgi intermediate compartments. **(A-B)** Characterization of ARF4 dynamics. ARF4^EN^-Halo KI HeLa cells were stained with the HaloTag substrate JF_552_-CA. **(A)** Imaging of ARF4^EN^-Halo cells with a spinning disk confocal microscope highlights three types of events (shown by black arrows in each crop): Ia) tubular connections between peripheral structures, Ib) tubular connections between peripheral structures and Golgi-associated structures, II) translocation of a peripheral structure to become a GA-ERGIC. Images were acquired at 2.4 frames per second. Scale bar 10 µm and 1 µm in the crops. **(B)** Frequency of the three types of ARF4 dynamic events calculated using confocal time-lapses acquired at 0.5 frames per second. n = 9. n = number of cells from three independent biological replicates. Error bars represent mean and SD. **(C-D)** Quantification of the instantaneous velocity of ARF4 structures. **(C)** Scatter dot plot shows the median instantaneous velocity from all trajectories from each cell (black dots). Error bars represent mean and SEM. Paired t test (**, P < 0.05, n = 7, n = number of cells from three independent biological replicates). **(D)** Relative frequency of the instantaneous velocity from all trajectories of elongated (grey bars) and rounded (black bars) ARF4 structures. **(E-G)** Characterization of ARF4-COPI dynamics. ARF4^EN^-Halo and βCOP^EN^-SNAP double KI HeLa cells were stained with the HaloTag substrate JF_571_-CA and the SNAP substrate JFX_650_-BG for STED microscopy (E and G) or JF_552_-CA and JFX_650_-BG for confocal microscopy (F). **(E)** STED image shows ARF4 tubular-vesicular structures (white arrows) at the Golgi (E i) and in the cell periphery (E ii) defined by COPI nanodomains (yellow arrows). Scale bar 5 µm and 1 µm in the crops. **(F)** Fast confocal time-lapse shows COPI (yellow arrows) associated to translocating ARF4 tubules (white arrows, Fi-ii). Images were acquired at 2.4 frames per second. Scale bar 10 µm and 2 µm in the crops. **(G)** STED time-lapse highlights the association of COPI to dynamic tubular-vesicular ARF4 structure. COPI clusters (yellow arrow) are observed at the neck of a membrane bridge connecting two peripheral structures (Ia) or a peripheral structure making a tubular connection (white arrows) with a GA structure (Ib). Tubular-vesicular structure tethering to the Golgi (white arrows, II). Images were acquired at 1 frame/5s. Scale bar 2 µm and 0.5 µm in the crops. **(H)** Schematic representation of the localization of COPI (magenta) on the ARF4 tubular-vesicular network and the dynamic events observed. GA-ERGIC=Golgi-associated ERGIC, p-ERGIC=peripheral ERGIC, CA=choloroalkene, BG=benzylguanine. Images were deconvolved, background subtracted and smoothed with a Gaussian filter.

As ARF4 ERGIC tubules defined by COPI were reminiscent of the tubular network described by Weigel and colleagues^10^, we tested the localization of endogenous ARF4 in respect to COPII machinery. The great majority of ARF4 p-ERGICs are associated to ERES punctae defined by endogenously SNAP-tagged Sec13 in both HeLa and RPE-1 cells (Fig. 2A-C). When analyzing the dynamics of ARF4 and COPII, ARF4 tubules are quickly remodeling around immobile ERES (Fig. 2D-F). Accordingly, ARF4 structures have a higher instantaneous velocity and diffusion coefficient than Sec13 structures (Fig. 2D-E, Extended Data Fig. 4A-B). Confocal time-lapse series show multiple different ARF4 tubules budding off from static ERES and connecting static ERES to one another in both double edited HeLa (Fig. 2F, Extended Data Fig. 4Ci-ii, Supplementary Movie 5) and RPE-1 cells (Extended Data Fig. 4D). Live-cell STED allowed us to follow the dynamic behavior of peripheral ERES near the Golgi and their relationship to GA-ERGICs (Fig. 2G). Interestingly, ARF4-positive ERGICs that are connected to stable ERES establish transient connections with GA-ERGICs (Fig. 2G, Supplementary Movie 6). GA-ERGICs are also directly connected to ERES as Sec13^EN^-SNAP punctae are seen associated to ARF4 GA-ERGICs in Hela and RPE-1 cells in live-cell STED images (Fig. 2H-I). To test whether ARF4 ERGICs are continuous with the ER, we expressed the soluble retention using selective hooks (RUSH)^23^ cargo TNFα-SBP-SNAP (TNFα-RUSH) in ARF4^EN^-Halo edited cells (Fig. 2J). Before the addition of biotin, TNFα-RUSH is retained in the ER by a Streptavidin-KDEL hook. Addition of biotin triggers release from the ER hook and trafficking out of the ER. Interestingly, without the addition of biotin, ER-resident TNFα-RUSH is able to freely diffuse into ARF4 p-ERGICs and GA-ERGICs, suggesting continuities between the lumen of the two organelles^10^. Taken together, we propose a model where ARF4 tubular-vesicular ERGIC structures are continuous with the ER and represent the entry gateway to the cis-Golgi via GA-ERGICs (Fig. 2K).

**Figure 2.**
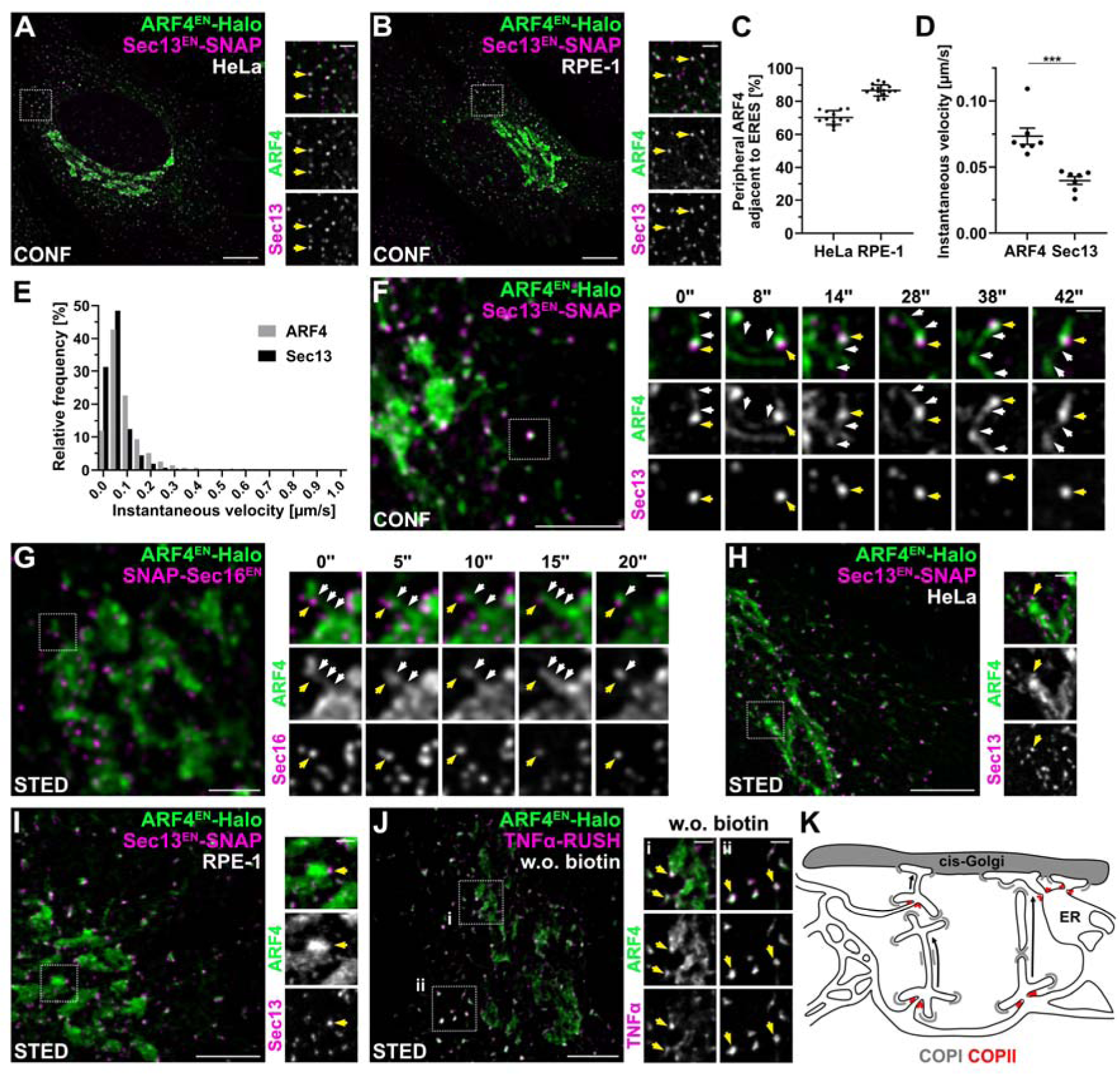
The ARF4 network dynamically remodels around static ERES. **(A-B)** Confocal images show peripheral ARF4 structures adjacent to ERES in HeLa and RPE-1 cells. ARF4^EN^-Halo and Sec13^EN^-SNAP double KI HeLa (A) and RPE-1 cells (B) were stained with the HaloTag substrate JF_552_-CA and the SNAP substrate JFX_650_-BG. ARF4 peripheral structures adjacent to ERES are highlighted by yellow arrows (A and B). Scale bar 10 µm and 2 µm in the crops. **(C)** Distribution of peripheral ARF4 structures adjacent to ERES in HeLa and RPE-1 cells. The percentage of peripheral ARF4 structures adjacent to ERES was calculated as described in the methods. HeLa n = 12, RPE-1 n = 15. n = number of cells from three independent biological replicates. Error bars represent mean and SD. **(D-G)** Characterization of ARF4-Sec13 dynamics. **(D-E)** Quantification of the instantaneous velocity of ARF4 and Sec13 structures. **(D)** Scatter dot plot shows the median instantaneous velocity from all trajectories from each cell (black dots). Error bars represent mean and SEM. Paired t test (***, P < 0.05, n = 7, n = number of cells from three independent biological replicates). **(E)** Relative frequency of the instantaneous velocity from all trajectories of ARF4 elongated and rounded structures (grey bars) and Sec13 punctae (black bars). **(F)** Confocal time-lapse imaging shows ARF4 tubular-vesicular structures dynamically remodeling around static ERES. ARF4^EN^-Halo and Sec13^EN^-SNAP double KI HeLa cells were stained with the Halo substrate JF_552_-CA and the SNAP substrate JFX_650_-BG. Images were acquired at 1 frame/2s. Scale bar 5 µm and 1 µm in the crops. **(G)** STED time-lapse imaging shows peripheral ARF4 establishing a tubular connection with a GA-ERGIC while still associated to ERES. ARF4^EN^-Halo and SNAP-Sec16^EN^ double KI HeLa cells were stained with the HaloTag substrate JF_571_-CA and the SNAP substrate JFX_650_-BG. Images were acquired at 1 frame/5s. Scale bar 2 µm and 0.5 µm in the crops. **(H and I)** STED imaging highlight ERES associated with GA-ERGICs in HeLa and RPE-1 cells. ARF4^EN^-Halo and Sec13^EN^-SNAP double KI HeLa (H) and RPE-1 cells (I) were stained with the HaloTag substrate JF_571_-CA and the SNAP substrate JFX_650_-BG. ERES associated to GA-ERGICs are highlighted by yellow arrows (H and I). Scale bar 5 µm and 1 µm in the crops. **(J)** STED imaging of TNFα-SBP-SNAP (TNFα-RUSH) in ARF4^EN^-Halo HeLa cells shows accumulation of cargo (yellow arrows) in GA (i) and peripheral ARF4 ERGICs (ii) before the addition of biotin. ARF4^EN^-Halo cells were transfected with a plasmid encoding for TNFα-RUSH and stained with the HaloTag substrate JF_571_-CA and the SNAP substrate JFX_650_-BG. Scale bar 5 µm and 1 µm in the crops. **(K)** Schematic representation of the localization of ERES (red) and COPI (grey) on the ARF4 tubular-vesicular network and the dynamic events observed in light of published reports^9,10^. CA=choloroalkene, BG=benzylguanine. Images were deconvolved (F-J), background subtracted and smoothed with a Gaussian filter (A-B, F-J).

According to this proposed model, we would expect an anterograde cargo released from the ER to enter the ARF4 ERGIC network and fill p-ERGICs and GA-ERGICs before filling the cis-Golgi cisternae. To test this, we again took advantage of the RUSH system and transfected ARF4^EN^-Halo and ARF1^EN^-Halo cells with a plasmid encoding for a Transferrin receptor (TfR) transmembrane RUSH cargo chimera (TfR-SBP-SNAP, TfR-RUSH) (Fig. 3, Extended Data Fig. 5, Supplementary Movies 7-8). Since ARF1 is excluded from GA-ERGICs (Extended Data Fig. 2A-B), the ARF1^EN^-Halo KI serves here as a cis-Golgi marker (Extended Data Fig. 5). Before biotin addition, the cargo accumulates in ERES close to ARF4 GA-ERGICs (Fig. 3Ai) and p-ERGICs (Fig. 3Aii) in proximity to the cis-Golgi (Extended Data Fig. 5A). 5 minutes after addition of biotin, cargo is seen in ARF4-positive p-ERGICs and GA-ERGICs (Fig. 3Bi) and in ARF4-positive tubules (Fig. 3Bii). Cargo is observed at GA-ERGICs tightly tethered to the cis-Golgi defined by ARF1 (Extended Data Fig. 5B). After 10 minutes, cargo is still seen in transit through ARF4-positive tubules (Fig. 3Cii) but starts emptying out from ARF4 GA-ERGICs (Fig. 3Ci) and starts filling the cis-Golgi cisternae defined by ARF1 (Extended Data Fig. 5C). Quantification of the kinetics of TfR-RUSH arrival at the Golgi shows that the cargo fills GA-ERGICs (Fig. 3D) and peripheral tubular-vesicular structures (Fig. 3E-F) positive for ARF4 before arriving at the cis-Golgi. To be able to discern closely-juxtaposed structures like peripheral ERES from ARF4 p-ERGICs and GA-ERGICs from cis-Golgi cisternae, we performed live-cell STED RUSH assays (Fig. 3G-H). The enhancement in resolution provided by STED microscopy allows to also separate tubular-vesicular ARF4 compartments from closely associated COPII punctae. Prior to biotin addition, TfR-RUSH is observed in punctate structures close to ARF4 GA- and p-ERGICs, that represent ERES (Fig. 3G), and near ARF1-positive cis-Golgi cisternae (Fig. 3H). After biotin addition, TfR-RUSH fills GA-ERGICs tightly tethered to the cis-Golgi ARF1 cisterna (Fig. 3G-H). TfR tubular carriers, imaged upon release of aggregated FM4-TfR-SNAP from the ER, are also observed in cells where only βCOP has been endogenously tagged with Halo, excluding any possible artifacts due to ARF4 tagging (Fig. 3I). Remarkably, TfR localizes along the length of the tubules and is not enriched in COPI nanodomains (Fig. 3I, COPI indicated by yellow arrow). Taken together, our functional trafficking assays using using synchronized cargo release show that anterograde cargo percolates through a network of ARF4-positive tubular-vesicular ERGICs to reach GA-ERGICs before eventually entering the Golgi apparatus via yet unknown mechanisms.

**Figure 3.**
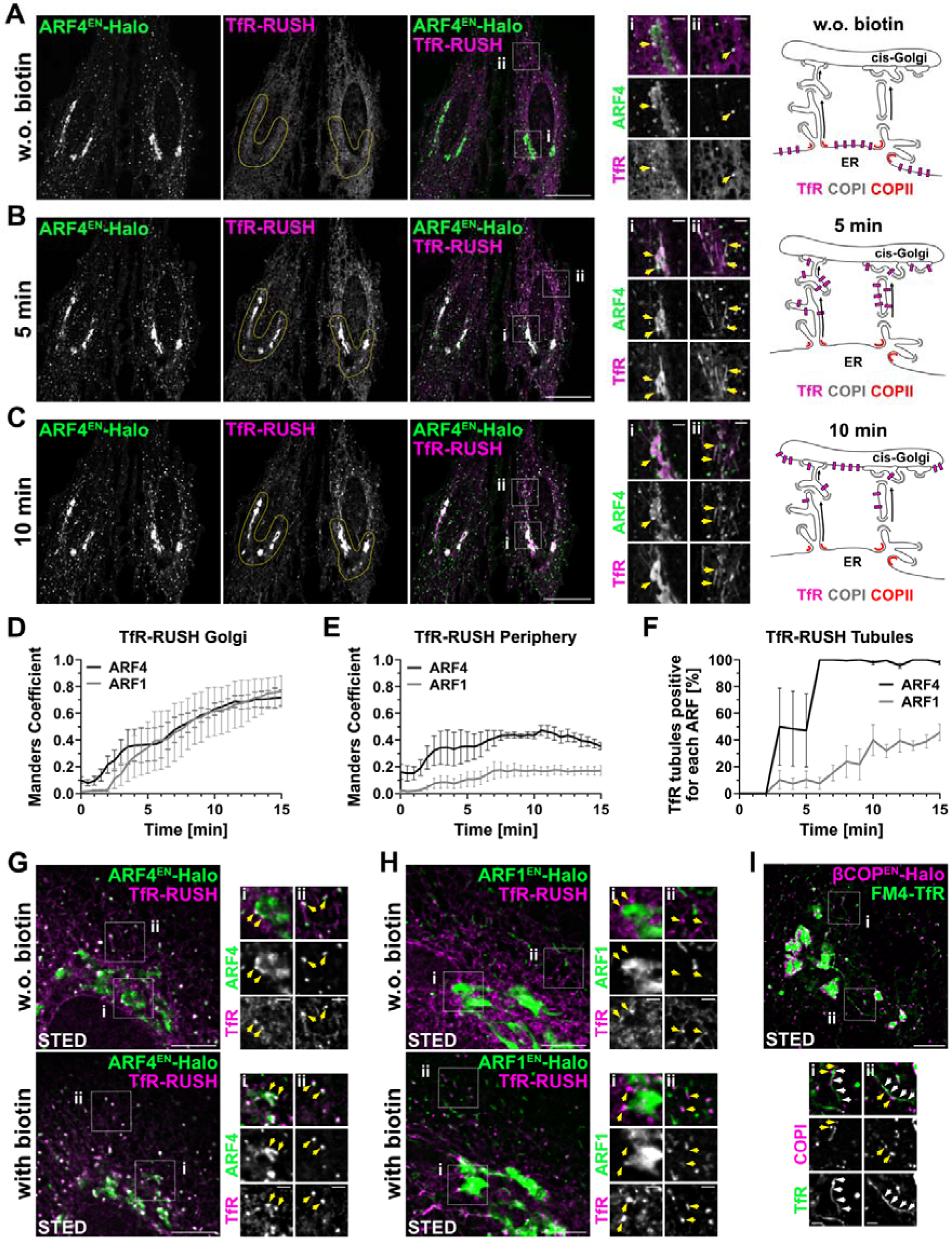
Anterograde cargo percolates through an ARF4 tubular network connecting ER and cis-Golgi. **(A-C)** Confocal time-lapse imaging of TfR-SBP-SNAP (TfR-RUSH) in ARF4^EN^-Halo HeLa cells reveals the path of anterograde cargoes from the ER to the Golgi. ARF4^EN^-Halo KI HeLa cells were transfected with a plasmid encoding for TfR-RUSH and stained with the HaloTag substrate JF_552_-CA and the SNAP substrate JFX_650_-BG. Images were acquired every 30 seconds. TfR-RUSH accumulation in the Golgi area is highlighted by yellow outlines in the overview images. TfR-RUSH accumulation in ARF4-positive GA-ERGICs and tubular-vesicular structures is indicated by yellow arrows in the crops. A schematic representation of the localization of TfR-RUSH cargo at different time points before and after biotin addition is provided. Scale bars are 10 µm and 1 µm in the crops. **(A)** TfR-RUSH before addition of biotin accumulates at the ER. **(B)** After addition of biotin, TfR-RUSH travels from the ER towards the Golgi via ARF4-positive tubular-vesicular structures. **(C)** TfR-RUSH filling the cis-Golgi while emptying out from GA-ERGICs. **(D-E)** Co-localization analysis (Manders coefficient) of TfR-RUSH in ARF4-(GA-ERGIC) or ARF1-(cis-Golgi) positive structures at the Golgi (D) and in the cell periphery (E) was calculated as described in the methods. **(F)** Percentage of TfR-RUSH tubules positive for ARF4 (ER-Golgi) or ARF1 (Golgi-ER or Golgi-PM) was calculated as described in the methods. Error bars represent SEM. ARF4 n = 4, ARF1 n = 4. n = number of independent RUSH assays (D-F). **(G-H)** STED imaging of TfR-RUSH in ARF4^EN^-Halo (E) and ARF1^EN^-Halo cells (F) highlights how TfR-RUSH cargo (yellow arrows) localizes to ARF4-positive GA-ERGICs tethered to cis-Golgi defined by ARF1. ARF4^EN^-Halo and ARF1^EN^-Halo KI HeLa cells were transfected with a plasmid encoding for TfR-RUSH and stained with the HaloTag substrate JF_571_-CA and the SNAP substrate JFX_650_-BG. Images were acquired before biotin addition and 10 mins after biotin addition. Scale bar 5 µm and 1 µm in the crops. **(I)** STED imaging of FM4-TfR-SNAP 5 mins after release from the ER with D/D solublizer in βCOP^EN^-Halo cells. TfR (white arrows) is not enriched in COPI nanodomains (yellow arrows). βCOP^EN^-Halo KI HeLa cells were transfected with a plasmid encoding for FM4-TfR-SNAP and stained with the HaloTag substrate JFX_650_-CA and the SNAP substrate JF_571_-BG. Trafficking assays were performed as described in the methods. CA=choloroalkene, BG=benzylguanine. Scale bar 5 µm and 1 µm in the crops. Images were deconvolved (E-G), background subtracted and smoothed with a Gaussian filter (A, E-G).

To test the role of ARF4 and COPI machinery on anterograde transport, we used a homozygous ARF4^EN^-Halo cell line^16^ and employed the Halo proteolysis targeting chimera (HaloPROTAC)^24^ strategy to acutely degrade endogenously Halo-tagged ARF4 (Fig. 4). Upon addition of HaloPROTAC, ARF4^EN^-Halo signal is strongly reduced by 90% as tested by labelling the remaining ARF4 chimera with a fluorescent ligand (Fig. 4A-B) and via Western Blot (Fig. 4C). Degradation of ARF4^EN^-Halo in a double KI cell line where also βCOP has been endogenously tagged with a SNAP tag (Fig. 4D), showed a marked reduction of membrane-associated βCOP (Fig. 4D-F). In particular, we observed less COPI coat associated to the Golgi (Fig. 4E) and to p-ERGICs (Fig. 4F), suggesting a role for ARF4 in the recruitment of COPI on GA- and p-ERGICs. Importantly, membrane recruitment of βCOP is restored upon overexpression of GFP-tagged ARF4 (Fig. 4C-F). In ARF4 KO cells^20^ (Extended Data Fig. 6), we could observe a less marked reduction of COPI signal only at the Golgi (Extended Data Fig. 6B), suggesting onset of compensatory effects. Supporting a role of ARF4 in the recruitment of COPI on ERGICs, we observe a significant increase in the recruitment of COPI on p-ERGICs in the ARF1 KO cells, where ARF4 is known to be strongly upregulated^20^ (Extended Data Fig. 6C). To investigate the effect on anterograde transport when ARF4 is acutely depleted, we performed TfR-RUSH assays in the absence or presence of HaloPROTAC (Fig. 4G-I, Supplementary Movies 9 and 10). Upon ARF4 depletion and subsequent reduced COPI recruitment to membranes, ER-Golgi transport of TfR-RUSH is accelerated as measured by the arrival of cargo in the perinuclear area (Fig. 4H). We also observed earlier onset of formation of TfR-positive tubules (Fig. 4I, Supplementary Movie 10), suggesting a regulatory role for ARF4 and COPI in anterograde ER-to-Golgi transport.

**Figure 4.**
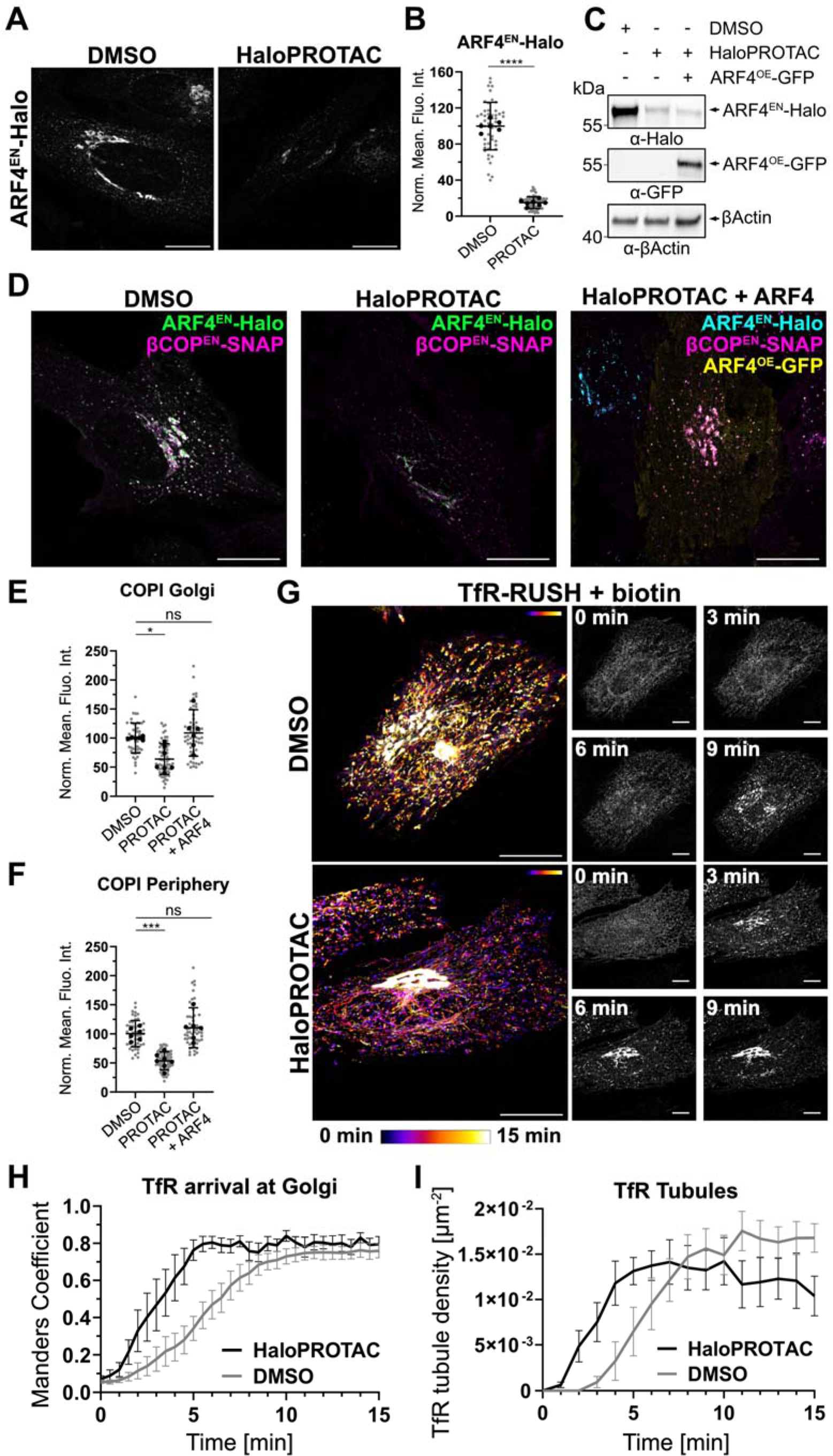
Acute degradation of ARF4 triggers COPI dissociation and mis-regulation of ER-Golgi transport. **(A)** ARF4^EN^-Halo HeLa cells were incubated with HaloPROTAC (1 µM, 24 h) or DMSO and stained with the HaloTag substrate JF_552_-CA. Scale bar 10 µm. **(B)** Quantification of the mean fluorescence intensity measured in ARF4^EN^-Halo HeLa cells treated with HaloPROTAC or DMSO. Values for individual cells are shown as grey dots and the mean value for each biological replicate as black dots. Error bars represent mean and SD. DMSO n = 6, HaloPROTAC n = 7, n = number of independent biological replicates (≥7 cells per replicate). Unpaired t test (****, P < 0.05). **(C)** Immunoblot to detect ARF4^EN^-Halo or ARF4^OE^-GFP fusion proteins in ARF4^EN^-Halo KI HeLa lysate with anti-Halo antibodies, anti-GFP antibodies and anti-βActin primary antibody as loading control. Cells were treated with DMSO or HaloPROTAC as indicated in the figure. **(D-F)** Acute degradation of ARF4^EN^-Halo with HaloPROTAC reduces COPI recruitment to GA- and p-ERGIC membranes. **(D)** ARF4^EN^-Halo and BetaCOP^EN^-SNAP double KI HeLa cells were treated with DMSO or HaloPROTAC and stained with the HaloTag substrate JF_552_-CA to detect residual Halo fusions and the SNAP substrate JFX_650_-BG. Rescue experiments were carried out by transfecting cells with a plasmid encoding for ARF4^OE^-GFP. Scale bar 10 µm. **(E-F)** Quantification of the mean fluorescence intensity of COPI at the Golgi area (E) or in the cell periphery (F) measured in ARF4^EN^-Halo and BetaCOP^EN^-SNAP double KI HeLa cells treated with DMSO or HaloPROTAC. Values for individual cells are shown as grey dots and the mean value for each biological replicate as black dots. Error bars represent mean and SD. DMSO n = 6, HaloPROTAC n = 7, HaloPROTAC+ARF4 = 6, n = number of independent biological replicates (≥7 cells per replicate). Ordinary one-way ANOVA versus DMSO (*,***, P < 0.05; ns = non-significant, P > 0.05). **(G-I)** Acute degradation of ARF4^EN^-Halo with HaloPROTAC affects ER-Golgi transport. **(G)** Temporal color map of ARF4^EN^-Halo KI HeLa cells transfected with a plasmid encoding for TfR-SBP-SNAP (TfR-RUSH) and treated with DMSO or HaloPROTAC. Cells were stained with the HaloTag substrate JF_552_-CA and the SNAP substrate JFX_650_-BG. Images were acquired every 30 seconds. Scale bar 10 µm. **(H)** Manders coefficient of TfR-RUSH at the Golgi was calculated as described in the methods. **(I)** TfR tubule density was calculated as described in the methods. Error bars represent SEM. DMSO n = 7, HaloPROTAC n = 6, n = number of independent RUSH assays (H-I). Norm. Mean Fluo. Int.= Normalized mean fluorescence intensity, CA=choloroalkene, BG=benzylguanine. Images were background subtracted and smoothed with a Gaussian filter.

## Discussion

The central vesicular dogma has accompanied the membrane trafficking field for the last 50 years, since the initial discoveries pioneered by Palade showing vesicles populating the cytoplasm of cells in early electron microscopy experiments^25^. COPI and COPII vesicles for transport to and from ERGIC compartments or the Golgi are often depicted in models explaining ER-Golgi transport. Recent advances in 3D correlative light and electron microscopy have afforded looking at the underlying membrane of ERES with unprecedented resolution and details^9,10^. These discoveries have brought a paradigm shift suggesting the presence of continuities between the ER and ERGIC membranes where cargoes would be channeled into a tubular-vesicular network via the action of COPII machinery in mammalian cells^9,10^. Transport via tubules rather than vesicles is supported by early observations of accumulation of cargoes into tubular trafficking intermediates upon release from the ER^2,26–29^. Here, we could dissect step-by-step how cargo is transferred from the ER to the Golgi and show a direct involvement of the small GTPase ARF4 and COPI machinery. We show that anterograde cargo transport between the ER and the Golgi is mediated by a dynamic interconnected network of tubules connecting the lumen of the ER with the lumen of p- and GA-ERGICs defined by the small GTPase ARF4 (Fig. 1 and Fig. 5). Super-resolution microcopy has been particularly instrumental in discerning the Golgi entry point via GA-ERGIC compartments (Fig. 2-3). It is unclear how cargo would then enter the first cis-Golgi cisterna. GA-ERGICs could be seen as pre-cisternae that could mature into a cis-Golgi cisterna for anterograde progression of cargoes. The static nature of these GA-ERGIC elements would otherwise suggest further transport from GA-ERGICs into the cis-Golgi via transport carriers or tubular connections. We speculate that loss of ARF4 and acquisition of ARF1 may drive the conversion switch between GA-ERGIC and cis-Golgi.

**Figure 5.**
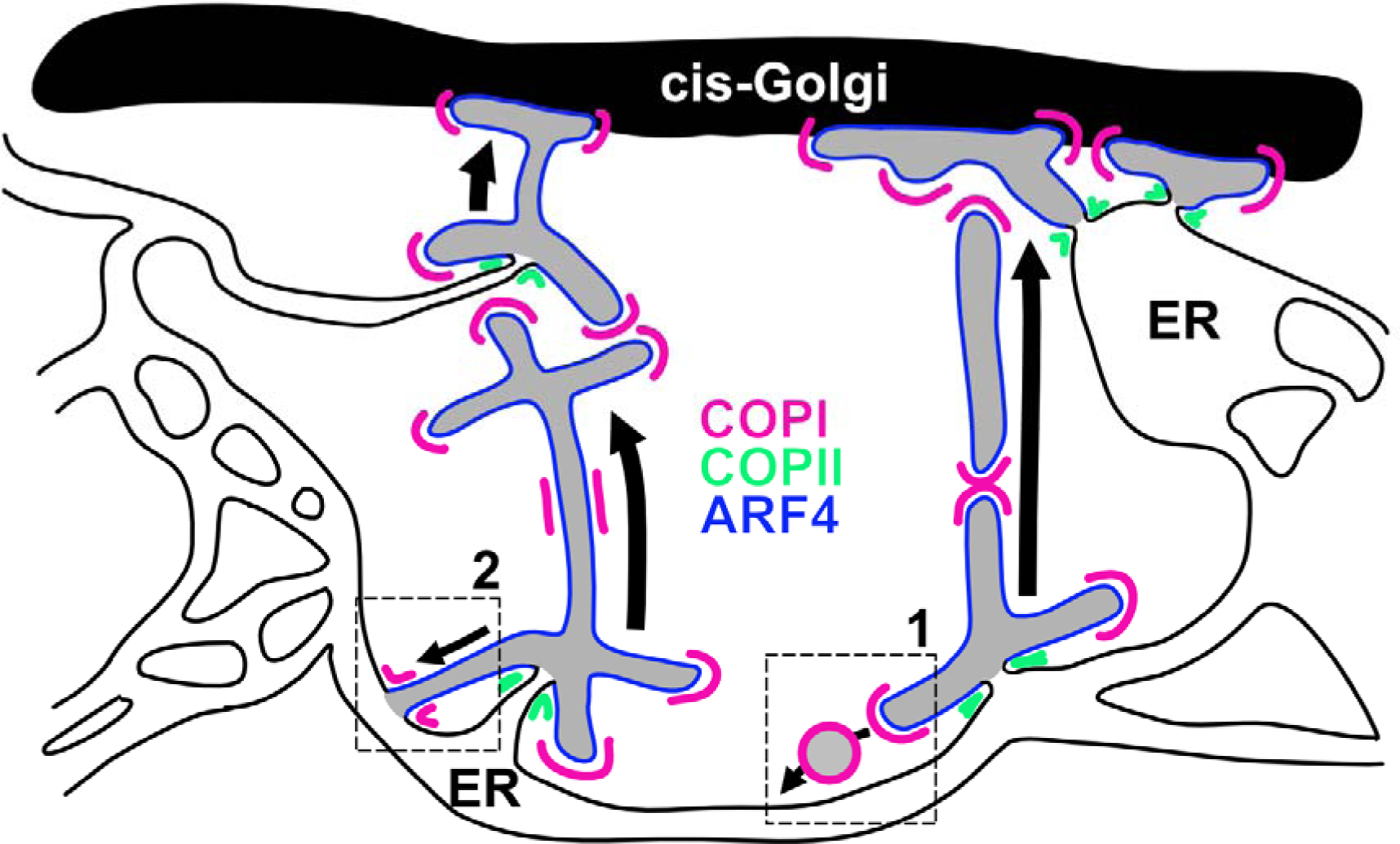
ER-to-Golgi transport occurs via a dynamic network of ARF4 tubules. ERGIC-to-ER retrieval could be mediated by either 1) COPI vesicles forming from ERGICs and fusing with the ER network below or 2) direct tubular connections with the ER.

But why would a network of tubules be better than vesicles? Vesicles are seen in yeast^30,31^. However, mammalian cells have a more complex endomembrane organization and different secretory and metabolic needs. Large tubular-vesicular structures would be more suitable for the transport of larger cargoes like collagen fibrils^32^, important for the deposition of the extracellular matrix. Additionally, ERES already associated to GA-ERGICs would provide a direct entry gateway to the Golgi without the need to be incorporated into small vesicular structures (Fig. 2H-I). In general, transport through a network of tubules would act as a “sieve” allowing time to separate anterograde-directed flow from ER residents and/or unfolded/aberrant proteins. Percolation through this tubular ERGIC ARF4 mesh would allow retrieval possibly via COPI-vesicles or ERGIC-to-ER continuities delivering retrograde cargoes to the ER network below where COPI could be speculated to act as a diffusion barrier (Fig. 5). In fact, COPI clusters are observed at the neck of tubular connections between ERGICs (Fig. 1G). Interestingly, anterograde cargo is distributed along the length of tubular carrier and is not enriched in COPI clusters (Fig. 3I). While ARF4 carriers move cargo forward, retrograde cargo could be sequestered away from the anterograde directed carriers by COPI. Further imaging technology advances pushing the axial resolution of volumetric light microscopy will be necessary to image such events, due to the overlapping geometry of ARF4 tubules and the ER network above/below. This ERGIC mesh model is well suited to explain the acceleration in anterograde transport caused by acute degradation of ARF4 and membrane dissociation of COPI (Fig. 4). ARF4-dependent recruitment of COPI on a tubular-vesicular ERGICs could act as a quality control brake, signaling that more material needs to be retrieved before tethering to the cis-Golgi could occur. The absence of ARF4/COPI could then lead to earlier and uncontrolled formation of ER-Golgi tubules.

Interestingly, ARF4 KO cells are defective in retrieval of ER KDEL clients^21^, suggesting that leakage outside of the cells could be due to accelerated ER-to-Golgi flow. However, analysis of the association of COPI in KO (Extended Data Fig. 6) vs acutely depleted cells (Fig. 4) show some discrepancies, suggesting that compensatory effects may be present in the KO cells. Additionally, fixed cells were analyzed^21^ and chemical fixation is known to lead to disruption of peripheral tubular-vesicular elements^16^. Strongly supporting our conclusions, ARF4 is known to be ∼4-fold upregulated in ARF1 KO cells^21^. Consequently, we observed a strong increase of COPI signal in p-ERGICs, due to the increase of ERGIC-bound ARF4 (Extended Data Fig. 6C). It is to be noted that upon depletion of ARF4, cargo filled tubules are still observed (Fig. 4). It is generally thought that ARFs could deform membranes and lead to formation of tubular-vesicular trafficking intermediates. However, this does not seem to be the case for ARF4 as tubules persist upon its depletion, suggesting alternative membrane bending mechanisms possibly via ARF paralogues (ARF1 or ARF5) or downstream recruitment of lipid modifying enzymes^19^.

Finally, we are just scratching the surface of the complexity of trafficking routes mediating ER-to-Golgi transport. Rab1 and the receptor SURF4 have been shown to regulate ER-Golgi transit via ERGIC53-negative tubular ERGIC compartments^33^. SURF4-dependent pathways would be particularly important in specific secretory cell types like insulin-secreting Beta cells^34^. However, Rab1 has also been shown to recruit GBF1^35^, the GDP-GTP exchange factor that would possibly trigger activation and recruitment of ARF4 on ERGICs, raising questions about the extent of the overlap between the various pathways.

## Materials and methods

### Cell culture

HeLa CCL-2 (ECACC General Collection) and Helaα^21^ were cultivated in DMEM (Gibco) supplemented with 10% FBS (Corning) and 1% penicillin/streptomycin (Thermo Fischer Scientific). hTERT RPE-1 (ATCC CRL-4000) were cultivated in DMEM-F12 supplemented with 10% FBS and 1% penicillin/streptomycin. All cells were grown at 37°C and 5% CO_2_.

HeLa cells were transfected using 2 µg of plasmid DNA and 6 µL of FuGENE (Promega) for gene editing or using a NEPA21 electroporation system (Nepa Gene) for trafficking assays. RPE-1 cells were transfected using the NEPA21 electroporation system. For electroporating HeLa or RPE-1 cells, 1 million cells were washed twice with Opti-MEM (Gibco) and resuspended in 90 μl Opti-MEM with 10 μg of DNA in an electroporation cuvette with a 2-mm gap. HeLa cells were electroporated with a poring pulse of 125 V, 3 ms pulse length, 50 ms pulse interval, 2 pulses, with decay rate of 10% and + polarity; consecutively with a transfer pulse of 20 V, 50 ms pulse length, 50 ms pulse interval, 5 pulses, with a decay rate of 40% and ± polarity. RPE-1 cells were electroporated with a poring pulse of 150 V, 5 ms pulse length, 50 ms pulse interval, 2 pulses, with decay rate of 10% and + polarity; consecutively with a transfer pulse of 20 V, 50 ms pulse length, 50 ms pulse interval, 5 pulses, with a decay rate of 40% and ± polarity. G418 (Gibco) was used at a concentration of 1 mg/mL for HeLa cells and 3 mg/mL for RPE-1 cells. Puromycin (Gibco) was used at concentration of 1 µg/mL for HeLa cells.

### Sample preparation for live-cell imaging

Imaging of live-cell samples was carried out on glass-bottom dishes (3.5 cm diameter, No. 1.5 glass; Cellvis) coated with fibronectin (Sigma-Aldrich). Live cells were labeled for 1 h using HaloTag and SNAP substrates at a concentration of 1 µM. To get rid of the excess of dye, cells were washed three times with growth medium after the staining and left for 1 h in an incubator at 37°C and 5% CO_2_. Imaging was done in FluoBrite DMEM (Gibco) supplemented with 10% FBS, 20 mM Hepes (Gibco), and 1× GlutaMAX (Gibco). Live-cell experiments were performed at 37°C. The combinations of Janelia Fluor dyes used in this study are listed in the figure legends^36–38^. For confocal imaging, we used JF_552_-CA^36^ and JFX_650_-BG^37^ to label the HaloTag and SNAP fusion proteins, respectively. For live-cell STED imaging, we replaced JF_552_-CA with JF_571_-CA^38^. The red-shifted spectra of the JF_571_ fluorophore allows efficient depletion using 775 nm, allowing facile two-color live-cell STED microscopy with a single depletion wavelength.

### Trafficking assays

RUSH assays were carried out by electroporating cells with 10 µg of plasmid DNA encoding for TfR-SBP-SNAP or TNFα-SBP-SNAP. Cells were seeded on glass-bottom dishes and incubated overnight at 37°C. Afterwards, cells were label as described above. Release of cargo from the ER was triggered by adding biotin (Sigma-Aldrich) to the live-cell imaging solution at a final concentration of 500 µM.

TfR-FM4 assays were carried out by electroporating cells with 10 µg of plasmid DNA encoding for TfR-FM4-SNAP. Cells were seeded on glass-bottom dishes and incubated overnight at 37°C. Afterwards, cells were label as described above. Release of TfR from the ER was triggered by adding D/D solubilizer (Takara Bio) to the live-cell imaging solution at a final concentration of 1 µM.

### Acute degradation with HaloPROTAC system

Acute degradation of endogenously Halo-tagged ARF4 was achieved using the HaloPROTAC-E compound [Dario Alessi and the MRC Reagents and Services division (University of Dundee)]. Cells were seeded in growth medium containing 1 µM HaloPROTAC and incubated for 24 h before imaging. Afterwards, cells were labeled as described above and imaged in live-cell imaging solution containing 1 µM HaloPROTAC.

### STED imaging

STED imaging was carried out on an Abberior Expert Line STED microscope supplied with 485, 561 and 646 nm laser lines and an Olympus Objective UPlanSApo 100x, NA 1.40, WD 0.13. The 775 nm depletion laser served to deplete both dyes in two-color STED measurements. Two-color images were acquired sequentially line by line. The detection was performed using avalanche photodiodes with detection windows set to 571-630 nm and 650-756 nm. Acquisition was done using the Instruments Development Team, Imspector Image Acquisition & Analysis Software v16.3.16118 (http://www.imspector.de). For confocal imaging, the pixel size was set to 60 nm or 80 nm for confocal time-lapses. For STED imaging, the pixel size was set to 30 nm or 40 nm for STED time-lapses. Live-cell STED and confocal imaging was carried out at 37°C.

### Spinning-disk confocal imaging

Spinning disk confocal imaging was carried out on a Nikon Spinning Disk Confocal CSU-W1 SoRa microscope equipped with 405, 445, 488, 515, 561, 594 and 638 nm laser lines and a Nikon Objective Plan Apo λ 60x oil, NA 1.4, WD 0.13. Two-color images were recorded by triggered acquisition using a Hamamatsu ORCA-Fusion digital sCMOS camera (2304×2304 pixels, 100 fps @ 2304×2048 pixels, 80% QE, 6.5 µm pixel size). Acquisition was performed with NIS-Elements AR Version 5.42.00 (https://www.microscope.healthcare.nikon.com/en_EU/products/software/nis-elements). The pixel size was 115 nm. Imaging was carried out at 37°C and 5% CO_2_.

### Image processing, quantification and statistical analysis

Confocal images were background subtracted and smoothed with a Gaussian filter with 1-pixel SD using ImageJ^39^. Raw STED and spinning disk confocal images were deconvolved using Richardson–Lucy deconvolution from the python microscopy PYME package (https://python-microscopy.org/), background subtracted and smoothed with a Gaussian filter with 1-pixel SD using ImageJ. All image analyses were performed using raw data unless otherwise stated. Data were plotted and statistically analyzed using GraphPad Prism.

The ARF4 dynamic events (Fig. 1B) were estimated using confocal time-lapse images acquired at 1 frame per second. The frequency of each event was counted, divided by the field of view and multiplied by 60 to obtain the number of events per µm² per min [µm^-2^min-^1^]. The percentage of peripheral ARF4 adjacent to ERES (Fig. 2C) was calculated by counting all ARF4-positive peripheral ERES punctae divided by the total amount of peripheral ERES punctae in ARF4^EN^-Halo and Sec13^EN^-SNAP double KIs HeLa and RPE-1 cells.

The Manders coefficient was calculated using background subtracted and smoothed confocal time-lapse images. To calculate the Manders coefficient of TfR-RUSH+ARF4/ARF1 at the Golgi area (Fig. 3C and Fig. 4H), a mask of the Golgi in the ARF4/ARF1 and TfR channels was determined by a thresholding method from ImageJ (Moments). In the HaloPROTAC treated cells (Fig. 4H), the amount of TfR localizing at the Golgi area was determined by the remaining ARF4^EN^-Halo signal. The Golgi masks of the TfR and ARF4/ARF1 channels were further analyzed with a Python script that calculated the Manders coefficient for the different time points of the time-lapses. The Manders coefficient of TfR-RUSH+ARF4/ARF1 in the cell periphery (Fig. 3D) was calculated by first excluding the Golgi area already measured as previously described. Afterwards, a mask of the structures in the cell periphery in the ARF4/ARF1 and TfR channels was determined by a thresholding method from ImageJ (MaxEntropy). The masks of the structures in the cell periphery of the TfR and ARF4/ARF1 channels were further analyzed with a Python script that calculated the Manders coefficient for the different time points of the time-lapses.

The percentage of TfR-RUSH tubules positive for ARF4 or ARF1 (Fig. 3E) was calculated by dividing the number of ARF4-or ARF1-positive TfR-RUSH tubules by the total number of TfR-RUSH tubules for each time point.

The mean fluorescence intensity of endogenous ARF4 (Fig. 4B) measured in HaloPROTAC-or DMSO-treated ARF4^EN^-Halo cells was measured in ImageJ using confocal images acquired with the same imaging parameters. Images were thresholded (Moments) and the mean fluorescence intensity in the Halo-PROTAC-treated cells was normalized to the mean fluorescence intensity of DMSO-treated cells. Data were statistically analyzed by an unpaired t test (Fig. 4B) using GraphPad Prism. P values are indicated in the figure legend. The mean fluorescence intensity of endogenous βCOP at the Golgi area or in the cell periphery measured in HaloPROTAC-or DMSO-treated ARF4^EN^-Halo cells (Fig. 4E-F) as well as in ARF4/ARF1 KO cells (Extended Data Fig. 6B-C) was measured in ImageJ using confocal images acquired with the same imaging parameters. The Golgi area selected for the measurement was determined by a thresholding method from ImageJ (Moments). To measure the mean fluorescence intensity in the cell periphery, the Golgi area already measured was excluded and peripheral structures were further thresholded (MaxEntropy). The mean fluorescence intensity of βCOP in the HaloPROTAC-treated cells and ARF4/ARF1 KO cells was normalized to the mean fluorescence intensity of βCOP in the DMSO-treated and WT cells, respectively. Data were statistically analyzed by ordinary one-way ANOVA (Fig. 3F-G and Extended Data Fig. 6B-C) using GraphPad Prism. P values are indicated in the figure legend.

Temporal projection maps (Fig. 4G) were generated using the Temporal-Color Code from stacks of time-lapse images in ImageJ.

The density of TfR-RUSH tubules measured in ARF4^EN^-Halo cells treated with HaloPROTAC or DMSO (Fig. 4I) was calculated by dividing the number of TfR-RUSH tubules by the area of the cell in nm^-2^ for each time point.

The tubule density measured in ARF4^EN^-Halo and ARF1^EN^-SNAP double KI HeLa cells was estimated using confocal time-lapse images acquired at 1 frame/2s (Extended Data Fig. 2D). The number of tubules positive for only ARF4, only ARF1 or for both was divided by the area of the cell and multiplied by 60 to get the tubule density per µm² per min [µm²/min].

### Characterization of ARF4-Sec13 dynamics

The ARF4- and Sec13-positive structures were tracked in space and time by using established software tools. For the ERES (Sec13 channel), which appear as generally round structures, the tracking was performed using TrackMate^40^. Spot detection with TrackMate was performed using the LoG (Laplacian of Gaussian) detector, an estimated object diameter of 0.6 µm, a quality threshold of 1, and subpixel localization was allowed. No further quality thresholding was applied. The tracking was performed with the LAP tracker, allowing a linking distance of 1.2 µm, a gap-closing max distance of 1.2 µm, and 1 gap-closing max frame gap.

However, TrackMate did not perform satisfactorily on ARF4-positive structures, due to their heterogeneous shape (from round structures to rather elongated tubular structures; Extended Data Fig. 3). To circumvent this, we leveraged the morphological resemblance between ARF4 structures and mitochondria and used MitoMeter, an algorithm for segmentation and tracking of elongated structures such as mitochondria^41^. MitoMeter segmentation was performed with a minimum structure area of 0.3 µm, and a maximum structure area of 200 µm. No custom Gaussian filtering and no threshold were applied. For the tracking, the maximum velocity threshold was set to 1. Fission and fusion events were considered. Further analysis of the tracking data was performed using a custom-made script coded in MATLAB (available on: https://github.com/jangulo4/Tracking_tubules). The ARF4-labeled perinuclear area was excluded from the analysis.

For each identified structure, we computed *(i)* the instantaneous velocity in every frame as the frame-by-frame displacement multiplied by the frame time and *(ii)* the diffusion coefficient (also referred to as instantaneous diffusion coefficient, D_1-4_), which was calculated from the time-averaged mean square displacement (TA-MSD). Specifically, for a particle trajectory with coordinate position vectors *xj*, sampled in time *N* times, the TA-MSD at time lag t_lag_ is calculated as:

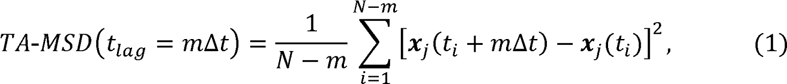

From here, the diffusion coefficient, D_l-4_, was computed by a linear fit of TA-MSo(t_lag_)= 4D_l-4_ t_lag_using the first four time lags, tlag = mΔt, for m= 1-4, where Δt is the frame time^42^. For these computations, we used only those trajectories with a minimum trajectory length of 12 frames.

Using the shape characterization provided by MitoMeter, the aspect ratio of the ARF4 structures was used for the classification of the diffusion parameters. The aspect ratio is defined as the major axis length divided by the minor axis length. Structures with an aspect ratio smaller than 1.6 are categorized as rounded, while those with a larger aspect ratio were considered elongated. When considering entire trajectories, the criterion to classify them as rounded or elongated was based on a majority rule, that is, if more than half of the frames were assigned to either of these categories, then the entire trajectory was assigned to that category. SuperPlots^43^ and histograms showing the diffusion parameters of ARF4 structures (Fig. 1C-D) or ARF4-Sec13 structures (Fig. 2D-E and Extended Data Fig. 4A-B) were plotted using GraphPad Prism. Paired t test (Fig. 1C, Fig. 2D, Extended Data Fig. 3D and Extended Data Fig. 4A) was performed with GraphPad Prism and p-values are indicated in the figure legend. The diffusion parameters together with the aspect ratio of ARF4 structures (Extended Data Fig. 3B-C) were plotted using GraphPad Prism.

### Plasmids for overexpression

TfR-FM4-Halo^44^ and YFP/SNAP-ERGIC53^16^ plasmids was previously described. To generate the RUSH plasmids Str-KDEL-TfR-SBP-SNAP and Str-KDEL-TNFalpha-SBP-SNAP, the original RUSH plasmids Str-KDEL-TfR-SBP-GFP and Str-KDEL-TNFα-SBP-GFP^23^, respectively, were linearized with SbfI and XbaI to cut out the GFP tag fragment and insert a SNAP tag fragment. To amplify the SNAP tag sequence out of the pSNAPf plasmid (New England Biolabs), the following primers were used: SbfI-SNAP-se: 5’-ATAACCTGCAGGTGACAAAGACTGCGAAATGAAG-3’ SNAP-XbaI-ase: 5’-TTAATCTAGATTAACCCAGCCCAGGCTTGCCCAG-3’.

To generate the ARF4-WT-GFP plasmid for overexpression, a pEGFP N3 (Clontech) vector was linearized with EcoRI and SalI to cut out the ARF1-WT fragment and the ARF4-WT fragment was inserted via Gibson assembly.

### CRISPR-Cas9 gene editing

The ARF4 genomic locus (gene ID 378) was targeted as previously described^16^. The ARF1 genomic locus (gene ID 375) was targeted as previously described^17^. The COPB1 genomic locus (gene ID 1315) was targeted as previously described^16^. The SEC13 genomic locus (gene ID 6396) was targeted with the guide RNA: 5’-ACGAGCAGTGACAAGACAGGTGG-3’ located after the stop codon in the SEC13 coding sequence. The SEC16A genomic locus (gene ID 9919) was targeted with the guide RNA: 5’-CCAGACGGGACCGTCTGGGGCGG-3’ located after the start codon in the SEC16A coding sequence. Guide RNAs designed using Benchling (https://www.benchling.com) did not have critical off-target matches when controlled with the CRISPR tool. The guide RNAs for ARF4, COPB1 and SEC13 were cloned by annealing oligoes and ligating them into the BpiI (Thermo Fisher Scientific) linearized vector pSpCas9(BB)-2A-Puro (pX459) V2.0^45^ (plasmid #62988; Addgene. Using the same strategy, the guide RNA oligoes to target the ARF1 and SEC16A locus were cloned into the vector SpCas9 pX330^46^ (plasmid #42230; Addgene).

The homology repair (HR) plasmids for ARF4-Halo and BetaCOP-Halo/SNAP were previously described^16^. The HR plasmid for ARF1-Halo/SNAP was cloned as previously described^17^. The HR plasmid for Sec13-SNAP was synthesized by GenScript with homology arms of ca. 1 kb each in a pUC57 vector backbone with BamHI and EcoRI sites to insert the desired tag a GS rich linker (GSGSGSGSGS). The HR plasmid for SNAP-Sec16 was synthesized by Twist Bioscience with homology arms of ca. 1 kb each in a pTWIST Amp High Copy plasmid (origin: pMB1) with HindIII, XhoI and SpeI sites for further downstream cloning. To avoid cutting by the Cas9 in HR plasmids, the protospacer adjacent motif site was mutagenized.

To generate the Sec13-SNAP-V5-PolyA-Puromycin^r^ HR plasmid, the empty Sec13 HR plasmid was linearized with BamHI and EcoRI. The BamHI-SNAP-V5-PolyA-Puromycin^r^-EcoRI fragment was cut out from a plasmid containing the tag and resistance cassette ^47^. To generate the LoxP-Puromycin^r^-LoxP-3xV5-SNAP-Sec16 HR plasmid, the empty Sec16 HR plasmid was linearized with HindIII and SpeI. The LoxP-Puromycin^r^-LoxP-3xV5-SNAP fragment was amplified out of the clathrin light chain CLC-SNAP HR plasmid described in^16^ with the following oligoes:

HindIII-LoxP-Puromycin^r^-se: 5’-ATGTCAAGCTTATAACTTCGTATAGCATACATTATACGAAGTTATCTGTGGAATGTGTGTCAGTTAGGGTGTGGAAAGTCCCCAGGCTCC-3’ SNAP-SpeI-ase: 5’-ACTGAACTAGTACCCAGCCCAGGCTTGCCCAGTCTG-3’.

The generation of gene edited ARF4^EN^-Halo, ARF1^EN^-Halo, BetaCOP^EN^-Halo, ARF4^EN^-Halo+ARF1^EN^-SNAP, ARF4^EN^-Halo+BetaCOP^EN^-SNAP in HeLa cells was previously described^16^. For the generation of the ARF4^EN^-Halo+Sec13^EN^-SNAP and ARF4^EN^-Halo+SNAP-Sec16^EN^ double KI in HeLa cells, Sec13-SNAP and SNAP-Sec16 HR plasmids and the respective guide RNA plasmid were transfected in the already existing ARF4^EN^-Halo cell line using FuGENE. For selection of positive clones, puromycin (1 µg/mL) was added to the cells 3 days after transfection. After selection and recovery, the resistance cassette flanked by LoxP sites (Sec16) was excised by transfection of 2 µg of pBS598 EF1alpha-EGFPcre plasmid (plasmid #11923; Addgene) that encodes for cre recombinase.

For the generation of gene edified ARF4^EN^-Halo, ARF4^EN^-Halo+ARF1^EN^-SNAP and ARF4^EN^-Halo+Sec13^EN^-SNAP in RPE-1 cells, HR plasmids and the respective guide RNA plasmids were transfected using a NEPA21 electroporation system (1 x 10^6^ cells, mixed with 5 µg of HR plasmid and 5 µg of guide RNA plasmid). Again, G418 (3 mg/mL) was added to the cells 3 days after transfection for selection of positive clones. To excise the resistance cassette flanked by LoxP sites (ARF4), cells were again electroporated with 10 µg of pBS598 EF1alpha-EGFPcre plasmid. Since the hTERT RPE-1 cell line already has a puromycin resistance cassette, the generation of double KI in this cell line was achieved by selecting only with G418. For the generation of the BetaCOP^EN^-Halo KI in WT, ARF4 KO and ARF1 KO Helaα cells, the BetaCOP HR plasmid and the guide RNA plasmid were transfected using FuGENE.

## Acknowledgments

This project was supported by the Deutsche Forschungsgemeinschaft (German Research Foundation) grants SFB958 (Project A25), SFB/TRR186 (Project A20). C.R. is supported by a Human Frontier Science Program (HFSP) early career award to F.B. We thank Stefan Donat and Jan Schmoranzer from the AMBIO imaging facility at Charité (Berlin) for the training on the spinning disk confocal microscope. We are grateful to Dario Alessi and the MRC Reagents and Services division (University of Dundee) for sharing the HaloPROTAC-E reagent. We thank Martin Spiess and lab for kindly providing ARF4 and ARF1 KO cells.

## Author contributions

L.W. and F.B. conceived the project. L.W., G.B., S.R., F.B. designed and performed experiments. L.W., G.B., S.R. and C.R. performed image analysis. S.E., S.H., P.L. generated plasmids and knock in cell lines. J.G. and L.D.L. synthesized imaging probes and provided technical expertise. J.A. and F.C. performed the tracking analysis. L.W. and F.B. wrote the manuscript with input from all authors.

## References

1 Aridor, M. A tango for coats and membranes: New insights into ER-to-Golgi traffic. Cell Rep 38 (2022). 10.1016/j.celrep.2021.110258

2 Bannykh, S. I., Rowe, T. & Balch, W. E. The organization of endoplasmic reticulum export complexes. J Cell Biol 135, 19–35 (1996). 10.1083/jcb.135.1.19

3 Miller, E. A. & Schekman, R. COPII - a flexible vesicle formation system. Curr Opin Cell Biol 25, 420–427 (2013). 10.1016/j.ceb.2013.04.005

4 Lujan, P., Angulo-Capel, J., Chabanon, M. & Campelo, F. Interorganelle communication and membrane shaping in the early secretory pathway. Curr Opin Cell Biol 71, 95–102 (2021). 10.1016/j.ceb.2021.01.010

5 Appenzeller-Herzog, C. & Hauri, H. P. The ER-Golgi intermediate compartment (ERGIC): in search of its identity and function. J Cell Sci 119, 2173–2183 (2006). 10.1242/jcs.03019

6 Pepperkok, R. et al. Beta-Cop Is Essential for Biosynthetic Membrane-Transport from the Endoplasmic-Reticulum to the Golgi-Complex in-Vivo. Cell 74, 71–82 (1993). doi: 10.1016/0092-8674(93)90295-2

7 Scales, S. J., Pepperkok, R. & Kreis, T. E. Visualization of ER-to-Golgi transport in living cells reveals a sequential mode of action for COPII and COPI. Cell 90, 1137–1148 (1997). 10.1016/s0092-8674(00)80379-7

8 Stephens, D. J., Lin-Marq, N., Pagano, A., Pepperkok, R. & Paccaud, J. P. COPI-coated ER-to-Golgi transport complexes segregate from COPII in close proximity to ER exit sites. J Cell Sci 113 (Pt 12), 2177–2185 (2000). 10.1242/jcs.113.12.2177

9 Shomron, O. et al. COPII collar defines the boundary between ER and ER exit site and does not coat cargo containers. J Cell Biol 220 (2021). 10.1083/jcb.201907224

10 Weigel, A. V. et al. ER-to-Golgi protein delivery through an interwoven, tubular network extending from ER. Cell 184, 2412-+ (2021). 10.1016/j.cell.2021.03.035

11 Raote, I. et al. TANGO1 builds a machine for collagen export by recruiting and spatially organizing COPII, tethers and membranes. Elife 7 (2018). 10.7554/eLife.32723

12 Raote, I. et al. A physical mechanism of TANGO1-mediated bulky cargo export. Elife 9 (2020). 10.7554/eLife.59426

13 Santos, A. J., Raote, I., Scarpa, M., Brouwers, N. & Malhotra, V. TANGO1 recruits ERGIC membranes to the endoplasmic reticulum for procollagen export. Elife 4 (2015). 10.7554/eLife.10982

14 Adarska, P., Wong-Dilworth, L. & Bottanelli, F. ARF GTPases and Their Ubiquitous Role in Intracellular Trafficking Beyond the Golgi. Front Cell Dev Biol 9, 679046 (2021). 10.3389/fcell.2021.679046

15 Popoff, V. et al. Several ADP-ribosylation Factor (Arf) Isoforms Support COPI Vesicle Formation. Journal of Biological Chemistry 286, 35634–35642 (2011). 10.1074/jbc.M111.261800

16 Wong-Dilworth, L. et al. STED imaging of endogenously tagged ARF GTPases reveals their distinct nanoscale localizations. Journal of Cell Biology (2023). 10.1083/jcb.202205107

17 Bottanelli, F. et al. A novel physiological role for ARF1 in the formation of bi-directional tubules from the Golgi. Mol Biol Cell (2017). 10.1091/mbc.E16-12-0863

18 Chun, J., Shapovalova, Z., Dejgaard, S. Y., Presley, J. F. & Melancon, P. Characterization of class I and II ADP-ribosylation factors (Arfs) in live cells: GDP-bound class II Arfs associate with the ER-Golgi intermediate compartment independently of GBF1. Mol Biol Cell 19, 3488–3500 (2008). 10.1091/mbc.E08-04-0373

19 Ben-Tekaya, H., Kahn, R. A. & Hauri, H. P. ADP ribosylation factors 1 and 4 and group VIA phospholipase A(2) regulate morphology and intraorganellar traffic in the endoplasmic reticulum-Golgi intermediate compartment. Mol Biol Cell 21, 4130–4140 (2010). 10.1091/mbc.E10-01-0022

20 Volpicelli-Daley, L. A., Li, Y. W., Zhang, C. J. & Kahn, R. A. Isoform-selective effects of the depletion of ADP-ribosylation factors 1-5 on membrane traffic. Mol Biol Cell 16, 4495–4508 (2005). 10.1091/mbc.E04-12-1042

21 Pennauer, M., Buczak, K., Prescianotto-Baschong, C. & Spiess, M. Shared and specific functions of Arfs 1-5 at the Golgi revealed by systematic knockouts. J Cell Biol 221 (2022). 10.1083/jcb.202106100

22 Bottanelli, F. et al. Two-colour live-cell nanoscale imaging of intracellular targets. Nat Commun 7, 10778 (2016). 10.1038/ncomms10778

23 Boncompain, G. et al. Synchronization of secretory protein traffic in populations of cells. Nat Methods 9, 493–498 (2012). 10.1038/nmeth.1928

24 Tovell, H. et al. Rapid and Reversible Knockdown of Endogenously Tagged Endosomal Proteins via an Optimized HaloPROTAC Degrader. ACS Chem Biol 14, 882–892 (2019). 10.1021/acschembio.8b01016

25 Palade, G. Intracellular aspects of the process of protein synthesis. Science 189, 867 (1975). 10.1126/science.189.4206.867-b

26 Presley, J. F. et al. ER-to-Golgi transport visualized in living cells. Nature 389, 81–85 (1997). 10.1038/38001

27 Aridor, M. et al. The Sar1 GTPase coordinates biosynthetic cargo selection with endoplasmic reticulum export site assembly. J Cell Biol 152, 213–229 (2001). 10.1083/jcb.152.1.213

28 Ben-Tekaya, H., Miura, K., Pepperkok, R. & Hauri, H. P. Live imaging of bidirectional traffic from the ERGIC. Journal of cell science 118, 357–367 (2005). 10.1242/jcs.01615

29 Simpson, J. C., Nilsson, T. & Pepperkok, R. Biogenesis of tubular ER-to-Golgi transport intermediates. Mol Biol Cell 17, 723–737 (2006). 10.1091/mbc.e05-06-0580

30 Barlowe, C. et al. Copii - a Membrane Coat Formed by Sec Proteins That Drive Vesicle Budding from the Endoplasmic-Reticulum. Cell 77, 895–907 (1994). Doi 10.1016/0092-8674(94)90138-4

31 Melero, A., Boulanger, J., Kukulski, W. & Miller, E. A. Ultrastructure of COPII vesicle formation in yeast characterized by correlative light and electron microscopy. Mol Biol Cell 33, ar122 (2022). 10.1091/mbc.E22-03-0103

32 Raote, I. & Malhotra, V. Protein transport by vesicles and tunnels. J Cell Biol 218, 737–739 (2019). 10.1083/jcb.201811073

33 Yan, R., Chen, K., Wang, B. & Xu, K. SURF4-induced tubular ERGIC selectively expedites ER-to-Golgi transport. Dev Cell 57, 512–525 e518 (2022). 10.1016/j.devcel.2021.12.018

34 Saegusa, K. et al. Cargo receptor Surf4 regulates endoplasmic reticulum export of proinsulin in pancreatic beta-cells. Commun Biol 5, 458 (2022). 10.1038/s42003-022-03417-6

35 Monetta, P., Slavin, I., Romero, N. & Alvarez, C. Rab1b interacts with GBF1 and modulates both ARF1 dynamics and COPI association. Mol Biol Cell 18, 2400–2410 (2007). 10.1091/mbc.E06-11-1005

36 Zheng, Q. et al. Rational Design of Fluorogenic and Spontaneously Blinking Labels for Super-Resolution Imaging. ACS Cent Sci 5, 1602–1613 (2019). 10.1021/acscentsci.9b00676

37 Grimm, J. B. et al. A General Method to Improve Fluorophores Using Deuterated Auxochromes. Jacs Au 1, 690–696 (2021). 10.1021/jacsau.1c00006

38 Grimm, J. B. et al. A general method to fine-tune fluorophores for live-cell and in vivo imaging. Nat Methods 14, 987–994 (2017). 10.1038/nmeth.4403

39 Abramoff, M. D., Magalhaes, P.J., Ram, S.J. Image Processing with ImageJ. Biophotonics International 11, 36–42 (2004).

40 Tinevez, J. Y. et al. TrackMate: An open and extensible platform for single-particle tracking. Methods 115, 80–90 (2017). 10.1016/j.ymeth.2016.09.016

41 Lefebvre, A., Ma, D., Kessenbrock, K., Lawson, D. A. & Digman, M. A. Automated segmentation and tracking of mitochondria in live-cell time-lapse images. Nat Methods 18, 1091–1102 (2021). 10.1038/s41592-021-01234-z

42 Manzo, C. & Garcia-Parajo, M. F. A review of progress in single particle tracking: from methods to biophysical insights. Rep Prog Phys 78, 124601 (2015). 10.1088/0034-4885/78/12/124601

43 Lord, S. J., Velle, K. B., Mullins, R. D. & Fritz-Laylin, L. K. SuperPlots: Communicating reproducibility and variability in cell biology. J Cell Biol 219 (2020). 10.1083/jcb.202001064

44 Bottanelli, F. & Schroeder, L. Stimulated Emission Depletion (STED) Imaging of Clathrin-Mediated Endocytosis in Living Cells. Methods Mol Biol 1847, 189–195 (2018). 10.1007/978-1-4939-8719-1_14

45 Ran, F. A. et al. Genome engineering using the CRISPR-Cas9 system. Nat Protoc 8, 2281–2308 (2013). 10.1038/nprot.2013.143

46 Cong, L. et al. Multiplex genome engineering using CRISPR/Cas systems. Science 339, 819–823 (2013). 10.1126/science.1231143

47 Stockhammer, A. et al. When less is more – Endogenous tagging with TurboID as a tool to study the native interactome of adaptor protein complexes. BioRxiv (2022). doi: 10.1101/2021.11.19.469212

