## Extended Data files for "Nanoscale imaging reveals the mechanisms of ER-to-Golgi transport via a dynamic tubular-vesicular network"

**Extended Data Figure 1. ARF4 tubular-vesicular structures are defined by the ERGIC marker ERGIC53 in HeLa and RPE-1 cells. (A-D)** Confocal (A-B) and STED (C-D) images show that ARF4 localizes to GA-ERGICs (A-D, i) and p-ERGICs (A-D, ii) defined by the ERGIC marker ERGIC53 in HeLa (A and C) and RPE-1 cells (B and D). HeLa and RPE-1 ARF4^EN^-Halo KI cells were transfected with a plasmid encoding for either YFP-ERGIC53 or SNAP-ERGIC53 as indicated in the figure. Live cells were stained with the HaloTag substrate JFX_650_-CA (A-B) or JF_571_-CA (C-D) and the SNAP substrate JFX_650_-BG (C-D). Scale bar 10 µm and 2 µm in the crops (A-B) or 5 µm and 1 µm in the crops (C-D). GA-ERGIC=Golgi-associated ERGIC, p-ERGIC=peripheral ERGIC, CA=choloroalkene, BG=benzylguanine. Images were deconvolved (C-D), background subtracted and smoothed with a Gaussian filter (A-D).

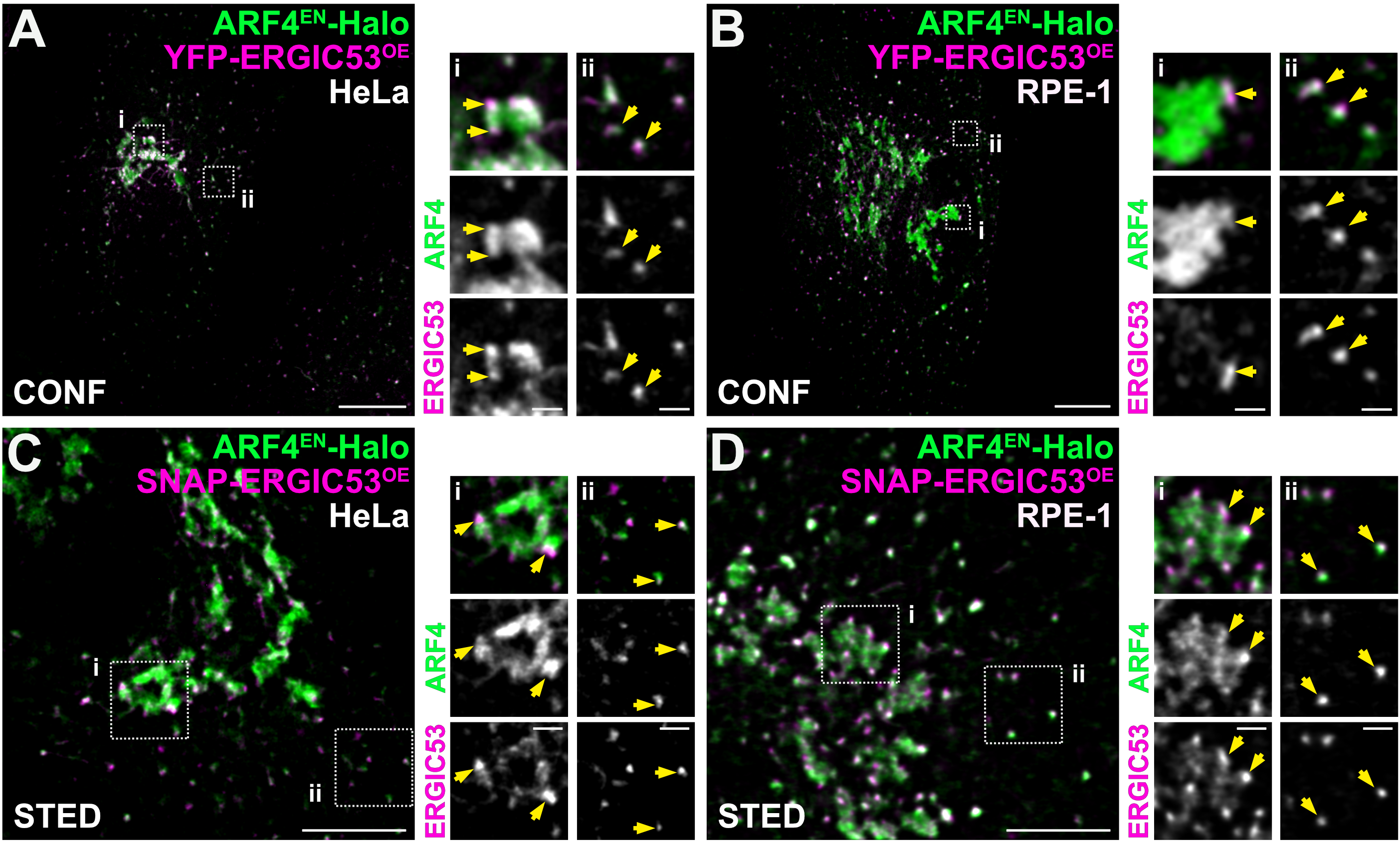

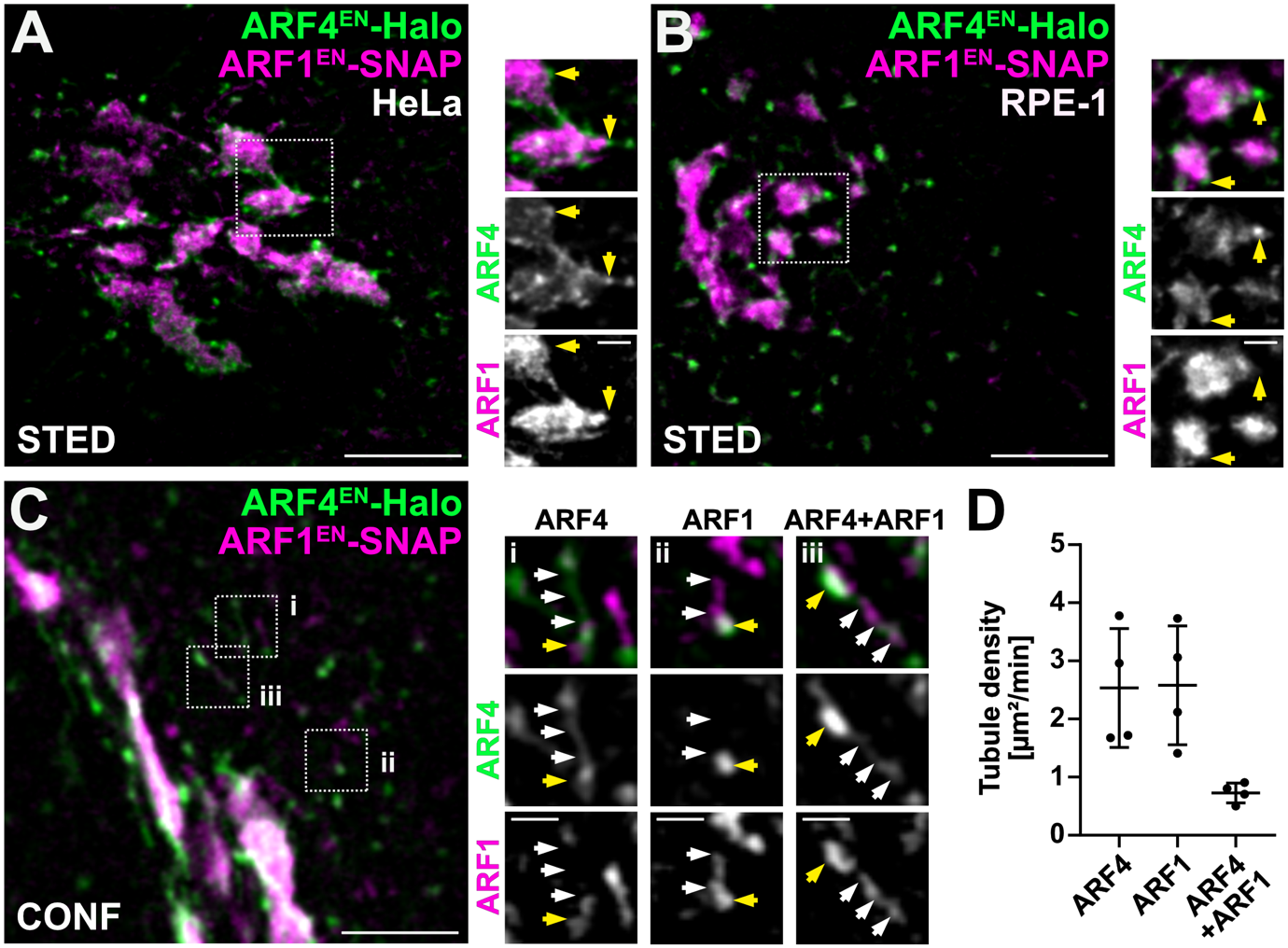
**Extended Data Figure 2. ARF4 defines GA-ERGICs tightly tethered to the cis-Golgi defined by ARF1 in HeLa and RPE-1 cells and sorting tubules segregated from ARF1 Golgi-to-ER retrograde tubules. (A-B)** ARF4^EN^-Halo and ARF1^EN^-SNAP double KI HeLa (A) and RPE-1 cells (B) were stained with the HaloTag substrate JF_571_-CA and the SNAP substrate JFX_650_-BG. GA-ERGICs are highlighted by yellow arrows. Scale bar 5 µm and 1 µm in the crops. **(C)** Confocal imaging shows that ARF4 sorting tubules (i) are segregated from ARF1 Golgi-to-ER retrograde tubules (ii). A subpopulation of tubules are positive for both ARFs (iii). Yellow arrows indicate p-ERGIC structures and white arrows indicate forming tubules. Scale bar 5 µm and 1 µm in the crops. **(D)** Tubule density was calculated using confocal time-lapses as described in the methods. Error bars represent mean and SD. n = 4, n = number of cells from three independent biological replicates. GA-ERGIC=Golgi-associated ERGIC, p-ERGIC=peripheral ERGIC, CA=choloroalkene, BG=benzylguanine. Images were deconvolded (A-B), background subtracted and smoothed with a Gaussian filter (A-C).

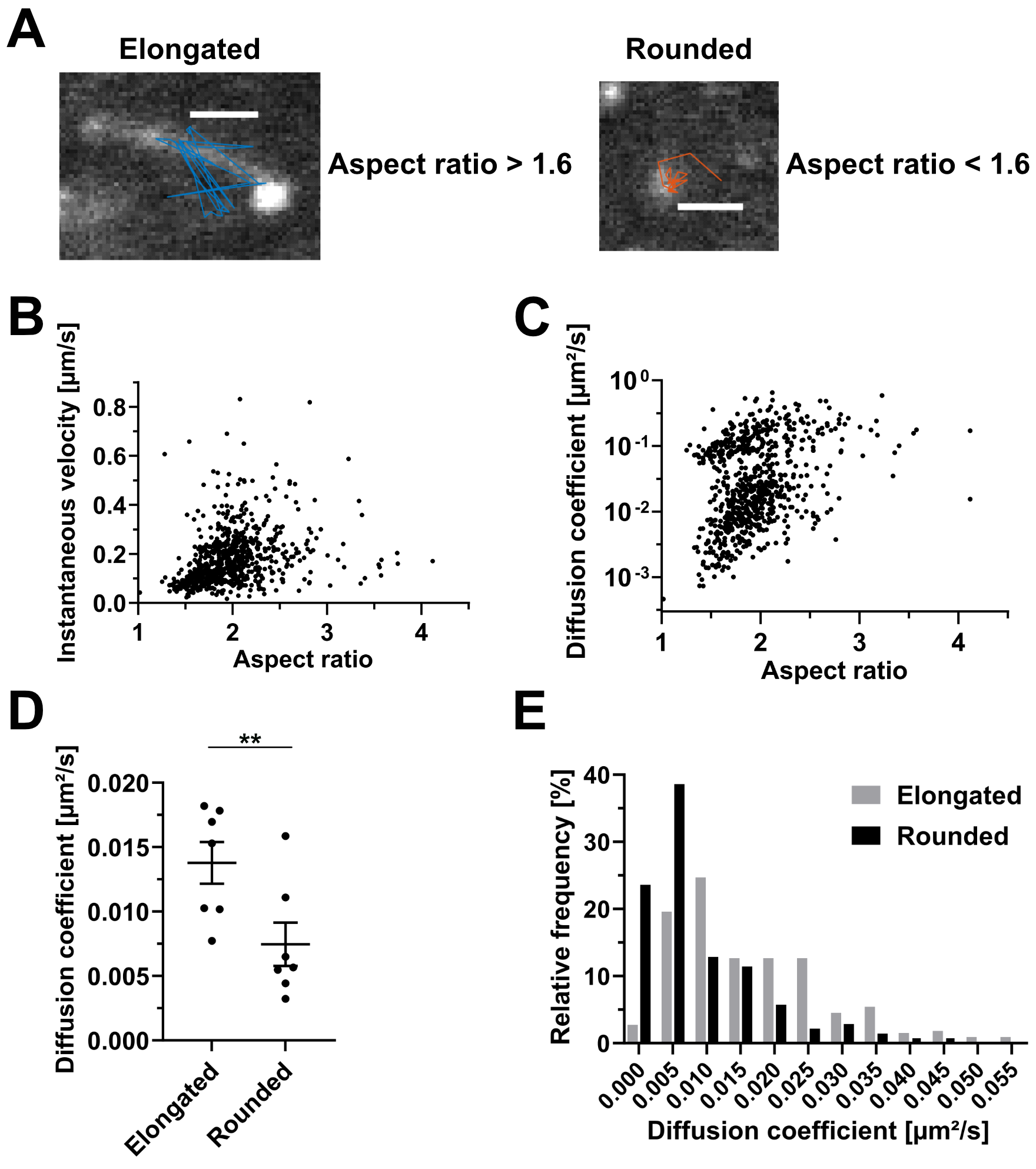

**Extended Data Figure 3. Characterization of ARF4 structures based on their mobility and shape. (A)** Classification of ARF4 structures. The aspect ratio was used to determine whether a structure is elongated or rounded. Structures with an aspect ratio above 1.6 are considered elongated and structures with an aspect ratio below 1.6 are considered rounded. The aspect ratio was determined as described in the methods. Scale bar 1 µm. **(B-C)** Dynamic properties of ARF4 structures. The instantaneous velocity (B) and diffusion coefficient (C) were calculated as described in the methods and plotted against the aspect ratio of each trajectory. **(D)** Quantification of the diffusion coefficient of ARF4 structures. Scatter dot plot shows the median diffusion coefficient from all trajectories from each cell (black dots). Error bars represent mean and SEM. Paired t test (**, P < 0.05, n = 7, n = number of cells from three independent biological replicates). **(E)** Relative frequency of the diffusion coefficient from all trajectories of elongated (grey bars) and rounded (black bars) ARF4 structures.

**Extended Data Figure 4. ARF4^EN^-Halo tubules dynamically remodel around static ERES in HeLa and RPE-1 cells. (A-B)** Quantification of the diffusion coefficient of ARF4 and Sec13 structures. **(A)** Scatter dot plot shows the median diffusion coefficient from all trajectories from each cell (black dots). Error bars represent mean and SEM. Paired t test (**, P < 0.05, n = 7, n = number of cells from three independent biological replicates). **(B)** Relative frequency of the diffusion coefficient from all trajectories of ARF4 elongated and rounded structures (grey bars) and Sec13 punctae (black bars). **(C-D)** ARF4^EN^-Halo and Sec13^EN^-SNAP double KI HeLa (C) and RPE-1 cells (D) were stained with the HaloTag substrate JF_552_-CA and the SNAP substrate JFX_650_-BG. White arrows highlight tubular structures defined by ARF4 and yellow arrows indicate static ERES defined by Sec13. **(C)** Confocal time-lapse imaging shows that peripheral ARF4 structures form tubular connections with other peripheral structures (i) and GA structures (ii) in HeLa cells. Crops from confocal time-lapse shown in Figure 2F. Scale bar 1 µm. **(D)** Confocal time-lapse imaging shows that peripheral ARF4 structures form and remodel around static ERES in RPE-1 cells (i). As observed for HeLa cells, tubules between p-ERGICs (ii) and GA-ERGICs (iii) are observed. Images were acquired at 1 frame/2s. Scale bar 5 µm and 1 µm in the crops. GA-ERGIC=Golgi-associated ERGIC, p-ERGIC=peripheral ERGIC, CA=choloroalkene, BG=benzylguanine. Images were background subtracted and smoothed with a Gaussian filter.

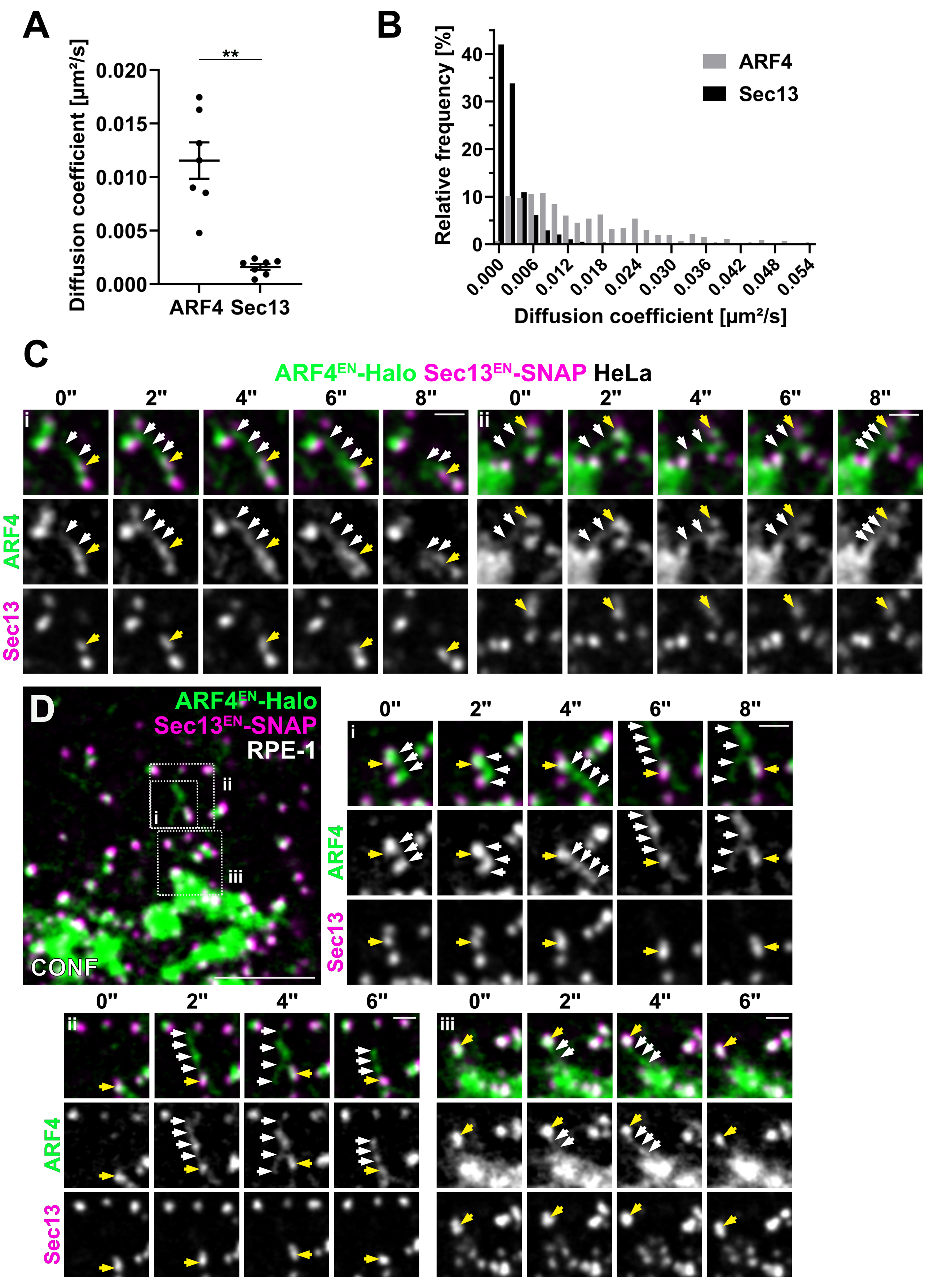

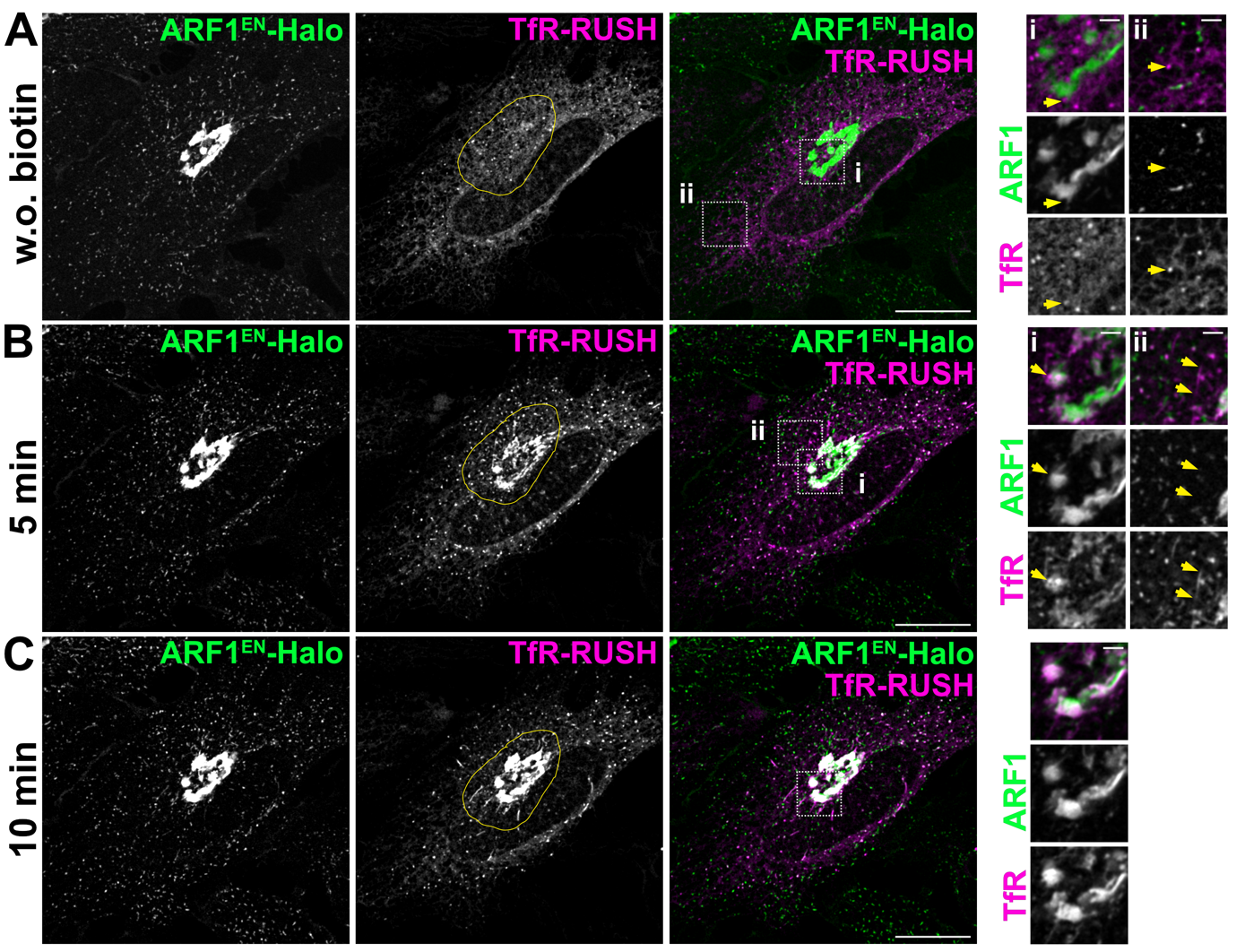

**Extended Data Figure 5. TfR-RUSH arrival at the cis-Golgi defined by the small GTPase ARF1.** ARF1^EN^-Halo KI HeLa cells were transfected with a plasmid encoding for TfR-SBP-SNAP (TfR-RUSH) and stained with the HaloTag substrate JF_552_-CA and the SNAP substrate JFX_650_-BG. Trafficking of TfR-RUSH was monitored before addition of biotin (A), after 5 (B) and 10 (C) minutes. Yellow outlines indicate TfR-RUSH accumulation at the cis-Golgi defined by ARF1. Yellow arrows in the crops highlight TfR-RUSH structures segregated from ARF1-positive structures and GA-ERGICs tethered to the cis-Golgi marked by ARF1. Images were taken every 30 seconds and were background subtracted and smoothed with a Gaussian filter. CA=choloroalkene, BG=benzylguanine. Scale bar 10 µm.

**Extended Data Figure 6. Effect of ARF4 and ARF1 KO on COPI recruitment. (A)** βCOP^EN^-Halo KI was created in WT, ARF4 KO and ARF1 KO HeLaα cells. Live cells were stained with the HaloTag substrate JFX_650_-CA. Images were background subtracted and smoothed with a Gaussian filter. Scale bar 10 µm. **(B-C)** Quantification of the mean fluorescence intensity of βCOP^EN^-Halo at the Golgi area (B) or in the cell periphery (C) measured in WT, ARF4 KO and ARF1 KO HeLaα cells. Values for individual cells are shown as grey dots and the mean value for each biological replicate as black dots. Error bars represent mean and SD. WT n = 8, ARF4 KO n = 6, ARF1 KO n= 6, n = number of independent biological replicates (≥10 cells per replicate). Ordinary one-way ANOVA versus WT (***,****, P < 0.05; ns = non-significant, P > 0.05). Images were background subtracted and smoothed with a Gaussian filter. CA=choloroalkene.

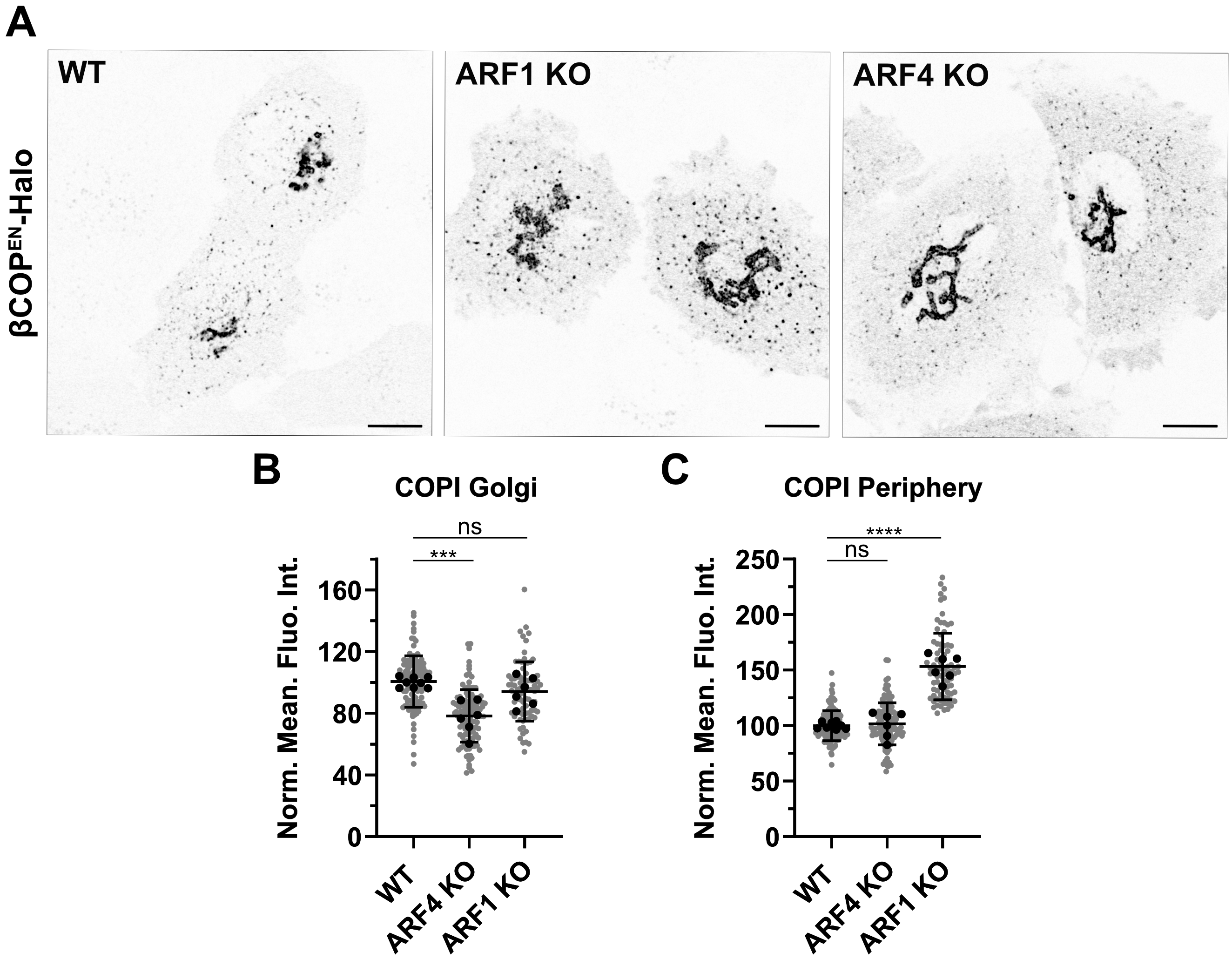

**Supplementary movies**

**Supplementary Movie 1. A highly pleiomorphic and quickly remodeling tubular-vesicular network positive for ARF4.** ARF4^EN^-Halo KI HeLa cells were stained with the HaloTag substrate JF_552_-CA and imaged with a spinning disk confocal microscope at 2.4 frames per second. Images were deconvolved, background subtracted and smoothed with a Gaussian filter. CA=choloroalkene. Scale bar 10 µm.

**Supplementary Movie 2. ARF4 tubules are mostly segregated from ARF1 tubules.** ARF4^EN^-Halo and ARF1^EN^-SNAP double KI HeLa cells were stained with the HaloTag substrate JF_552_-CA and the SNAP substrate JFX_650_-BG. Images were acquired at 1 frame/2s. Images were background subtracted and smoothed with a Gaussian filter. CA=choloroalkene, BG=benzylguanine. Scale bar 5 µm.

**Supplementary Movie 3. COPI remains associated to translocating ARF4 tubules.** ARF4^EN^-Halo and βCOP^EN^-SNAP double KI HeLa cells were stained with the HaloTag substrate JF_552_-CA and the SNAP substrate JFX_650_-BG. Images were acquired with a spinning disk confocal microscope at 2.4 frames per second. Images were deconvolved, background subtracted and smoothed with a Gaussian filter. CA=choloroalkene, BG=benzylguanine. Scale bar 10 µm.

**Supplementary Movie 4. ARF4 p-ERGICs establish tubular connections with GA-ERGICs and COPI localizes to the connecting tubular bridge.** ARF4^EN^-Halo and βCOP^EN^-SNAP double KI HeLa cells were stained with the HaloTag substrate JF_571_-CA and the SNAP substrate JFX_650_-BG. Images were acquired at 1 frame/5s. Images were deconvolved, background subtracted and smoothed with a Gaussian filter. CA=choloroalkene, BG=benzylguanine. Scale bar 2 µm.

**Supplementary Movie 5. ARF4 tubules remodel around immobile ERES in HeLa cells.** ARF4^EN^-Halo and Sec13^EN^-SNAP double KI HeLa cells were stained with the HaloTag substrate JF_552_-CA and the SNAP substrate JFX_650_-BG. Images were acquired at 1 frame/2s. Images were background subtracted and smoothed with a Gaussian filter. CA=choloroalkene, BG=benzylguanine. Scale bar 5 µm.

**Supplementary Movie 6. ARF4-positive p-ERGICs associated to ERES and establish transient tubular connections with GA-ERGICs.** ARF4^EN^-Halo and SNAP-Sec16^EN^ double KI HeLa cells were stained with the HaloTag substrate JF_571_-CA and the SNAP substrate JFX_650_-BG. Images were acquired at 1 frame/5s. Images were deconvolved, background subtracted and smoothed with a Gaussian filter. CA=choloroalkene, BG=benzylguanine. Scale bar 2 µm.

**Supplementary Movie 7. TfR-RUSH percolates through an ARF4 tubular-vesicular network.** ARF4^EN^-Halo KI HeLa cells were transfected with a plasmid encoding for TfR-RUSH and stained with the HaloTag substrate JF_552_-CA and the SNAP substrate JFX_650_-BG. Images were acquired every 30 seconds. Images were background subtracted and smoothed with a Gaussian filter. CA=choloroalkene, BG=benzylguanine. Scale bar 10 µm.

**Supplementary Movie 8. TfR-RUSH arrival at the cis-Golgi defined by ARF1.** ARF1^EN^-Halo KI HeLa cells were transfected with a plasmid encoding for TfR-RUSH and stained with the HaloTag substrate JF_552_-CA and the SNAP substrate JFX_650_-BG. Images were acquired every 30 seconds. Images were background subtracted and smoothed with a Gaussian filter. CA=choloroalkene, BG=benzylguanine. Scale bar 10 µm.

**Supplementary Movie 9. TfR-RUSH assay in the absence of HaloPROTAC.** ARF4^EN^-Halo KI HeLa cells were transfected with a plasmid encoding for TfR-RUSH and treated with DMSO as a control. Cells were stained with the SNAP substrate JFX_650_-BG. Images were acquired every 30 seconds. Images were background subtracted and smoothed with a Gaussian filter. BG=benzylguanine. Scale bar 10 µm.

**Supplementary Movie 10. TfR-RUSH assay in HaloPROTAC-treated ARF4 KI cells.** ARF4^EN^-Halo KI HeLa cells were transfected with a plasmid encoding for TfR-RUSH and treated with HaloPROTAC. Cells were stained with the SNAP substrate JFX_650_-BG. Images were acquired every 30 seconds. Images were background subtracted and smoothed with a Gaussian filter. BG=benzylguanine. Scale bar 10 µm.
